# Novel ‘housekeeping’ genes and an unusually heterogeneous distribution of transporter expression profiles in human tissues and cell lines, assessed using the Gini coefficient

**DOI:** 10.1101/155697

**Authors:** Steve O’Hagan, Marina Wright Muelas, Philip J. Day, Emma Lundberg, Douglas B. Kell

## Abstract

We analyse two comprehensive transcriptome datasets from human tissues and human-derived cell lines in terms of the expression profiles of the SLC and ABC families of membrane transporters. The Gini index (coefficient) characterises inequalities of distributions, and is used in a novel way to describe the distribution of the expression of each transporter among the different tissues and cell lines. In many cases, transporters exhibit extremely high Gini coefficients, even when their supposed substrates might be expected to be available to all tissues, indicating a much higher degree of specialisation than is usually assumed. This is consistent with divergent evolution from a more restricted set of ancestors. Similar trends hold true for the expression profiles of transporters in different cell lines, suggesting that cell lines exhibit largely similar transport behaviour to that of tissues. By contrast, the Gini coefficients for ABC transporters tend to be larger in cell lines than in tissues, implying that some kind of a selection process has taken place. In particular, with some exceptions such as olfactory receptors and genes involved in keratin production, transporter genes are significantly more heterogeneously expressed than are most non-transporter genes. The Gini index also allows us to determine those transcripts with the most stable expression; these often differ significantly from the ‘housekeeping’ genes commonly used for normalisation in transcriptomics and qPCR studies. The lowest four in tissues are FAM32A, ABCB7, MRPL21 and PCBP1, while the lowest three in cell lines are SF3B2, NXF1 and RBM45. PCBP1 is both reasonably highly expressed and has a low Gini coefficient in both tissues and cell lines, and is an excellent novel housekeeping gene. Overall, our analyses provide novel opportunities for the normalisation of genome-wide expression profiling data.

## Introduction

Given that the basic genome of a differentiated organism is constant between cells (and we here ignore epigenomics), what mainly discriminates one cell type from another is its expression profile. The ‘surfaceome’ – those proteins expressed on the cell surface – attracts our interest in particular, as it contains the transporters that determine which nutrients (and xenobiotics such as drugs) are taken up by specific cells (da Cunha et al., 2009; Palm and Thompson, 2017). Transporters are the second largest component of the membrane proteome (Almén et al., 2009), and also a (surprisingly) understudied clade (César-Razquin et al., 2015). They are classified into solute carriers (SLCs (Colas et al., 2016; Fredriksson et al., 2008; Hediger et al., 2013; Perland and Fredriksson, 2017; Schlessinger et al., 2010; Sreedharan et al., 2011)), mainly involved in uptake, and ABC transporters (ABCs), mainly involved in efflux (e.g. (Chen et al., 2016; Eadie et al., 2014; Montanari and Ecker, 2015; Rees et al., 2009)).

That transporters are also responsible for the uptake of pharmaceutical drugs and xenobiotics into cells, and their efflux therefrom (Colas et al., 2016; Dobson and Kell, 2008; Giacomini and Huang, 2013; Giacomini et al., 2010; Kell, 2015; Kell, 2016; Kell et al., 2013; Kell et al., 2011; Kell and Oliver, 2014; Lin et al., 2015; Stanley et al., 2009), means that to understand drug distributions we must understand transporter distributions. In many cases, we do not know either the ‘natural’ (O’Hagan and Kell, 2017c; Perland and Fredriksson, 2017) or the pharmaceutical drug substrates of these transporters, and one clue to this may be to understand their differential tissue distribution. Fortunately, we now have available a variety of expression profiles at the level of both the transcriptome (Fagerberg et al., 2014) and the proteome (Thul et al., 2017; Uhlén et al., 2015) that allow us to assess these. In the present work we used transcription profiles acquired as part of the tissue atlas (Uhlén et al., 2015) and cell atlas (Thul et al., 2017). Altogether there are four main datasets, namely 409 “SLCs” in 59 tissue types and 56 cell lines, and 48 ABCs in the same tissue types and cell lines. Some of the SLCs do not (yet) have the official terminology (Perland and Fredriksson, 2017; Perland et al., 2016; Sreedharan et al., 2011), but based on a variety of phylogenetic and other evidence, as well as their Uniprot annotations, they clearly have this function, and these are noted accordingly. In a similar vein, some of the ‘ABC’ families (especially family F) are probably not functionally membrane transporters, but they are nonetheless included.

## Materials and methods

Expression profiles were obtained as described (Thul et al., 2017), and the data extracted and extended in the form of Microsoft Excel sheets. mRNA sequencing was performed on Illumina HiSeq2000 and 2500 platforms (Illumina, San Diego, CA, USA) using the standard Illumina RNA-seq protocol with a read length of 2x100 bases. Transcript abundance estimation was performed using Kallisto (Bray et al., 2016) v0.42.4. For each gene, we report the abundance in ‘Transcripts Per Million’ (TPM) as the sum of the TPM values of all its protein-coding transcripts. For each cell line and tissue type, the average TPM values for replicate samples were used as abundance score. Thus each transcript level does represent an absolute value, but it is then normalised to the total expression in the particular sample.

Most of the analyses are self-explanatory. As in many of our cheminformatics analyses (e.g. (O’Hagan and Kell, 2017a; O’Hagan et al., 2015)) we used the KNIME software environment (Berthold et al., 2008; O’Hagan and Kell, 2017a; O’Hagan et al., 2015) (http://knime.org/), with visualisation often provided via the Tibco Spotfire™ software (Perkin-Elmer Informatics). A variety of means exist to capture variation (e.g. coefficient of variation, inter-quartile range, ratio of 20^th^:80^th^ percentiles); however, none of those captures the full range well, especially including the many zeroes (undetectable expression levels). The one that does is the Gini index, but it is somewhat unfamiliar; since it is a major focus here, we explain this next.

### The Gini index

The Gini index (Ceriani and Verme, 2012; Gini, 1909, 1912) or Gini coefficient (GC) is a non-parametric measure that is widely used in economics to describe the distribution of incomes between individuals in a given group or political jurisdiction (e.g. a country or region) (Kondo et al., 2012; Pickett and Wilkinson, 2015; Wilkinson and Pickett, 2009). As a summary statistic of the entire Lorenz curve (Lee, 1999) (and see Fig 1), it is effectively a statistical measure of the degree of variation represented in a set of values. It ranges between zero (no variation) and 1 (an extreme variation, in which all the non-zero values are contained in one individual or example), and is thereby easily understood. In economics by country, its recent values range from ca 0.25 (Slovenia) to ca 0.51 (Chile) http://data.worldbank.org/indicator/SI.POV.GINI. Clearly it can be used to describe the distribution of anything else, e.g. the structural diversity in chemical libraries (Weidlich and Filippov, 2016) (modulo the huge effects of encodings thereon (O’Hagan and Kell, 2017; Riniker and Landrum, 2013)), the sizes of individual plants (Damgaard and Weiner, 2000) or of their local productivity (Sadras and Bongiovanni, 2004) as a function of the total biomass in an ecosystem, or even the distribution of citations of bioinformatics papers (Wren, 2016). It has very occasionally been used in gene expression profiling studies (Ainali et al., 2012; Jiang et al., 2016; Torre et al., 2017; Tran, 2011). However, in each of these latter cases, the Gini index was used for choosing subsets of transcripts that differentiate rare cell types or diseases. Here we know the cell types and the novelty lies in using the Gini index to assess individual transporters in terms of the uniqueness of their expression levels. A more intuitive, graphical illustration of the derivation of the Gini coefficient is given in Fig 1.

**Figure 1.**
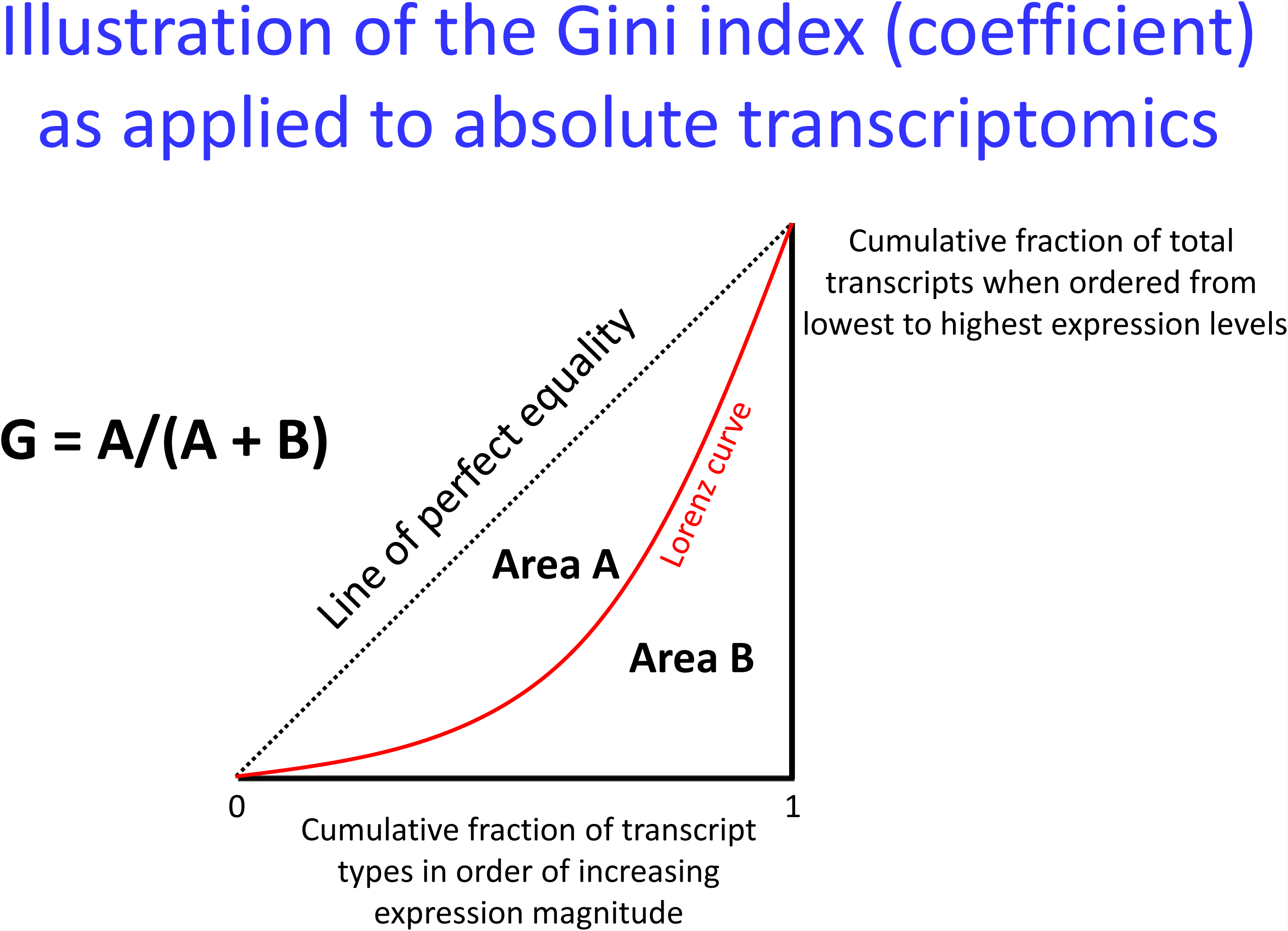
The Gini index. Many equivalent definitions are possible. In the usual form, the Gini coefficient is defined mathematically based on the Lorenz curve, which plots the proportion of the total income or wealth of a population (ordinate) that is earned cumulatively by the bottom x% of the population (see diagram) as x increases. Here ‘income’ is the percentage of total transcripts, while the ‘population’ is the individual transporter transcripts considered at one time. (The same general form results if the abscissa is reversed, starting with the top earners, where it takes on the appearance of the more familiar receiver-operator characteristic curve or ROC curve (Baker, 2003; Broadhurst and Kell, 2006; Linden, 2006).) The line at 45 degrees represents uniform expression of each transcript. The Gini coefficient can then be seen as the ratio of the area that lies between the line of equality and the Lorenz curve (labelled A in the figure) to the total area under the line of equality (labelled A and B), i.e., G = A / (A + B).

In the present work, the Gini Index was calculated using the **ineq** package (Achim Zeileis (2014). ineq: Measuring Inequality, Concentration, and Poverty. R package version 0.2-13. https://CRAN.R-project.org/package=ineq) in **R** (https://www.R-project.org/). These calculations were incorporated into KNIME via Knime’s R integration *R Snippet* node. The Rank Correlation used was Spearman’s rho, using the Knime *Rank Correlation node*.

### Immunohistochemistry

Immunohistochemical (IHC) images detailing protein expression patterns in 48 different normal tissues and 20 common cancer types are from the Human Protein Atlas database (www.proteinatlas.org). Tissue microarrays, immunostaining and image evaluation was performed as previously described (Uhlén et al., 2015). Briefly, 1mm duplicate cores were used for immunostaining using the following antibodies: HPA024575 for SLC22A12, HPA011885 for SLC6A18, HPA006539 for SLC2A14 (all from the Human Protein Atlas) and CAB037113 for PCBP1 (R1455 from Sigma-Aldrich). The immunostaining intensity and pattern was manually evaluated and scored.

## Results

### Variation in expression profiles of SLCs in tissues

Figure 2 shows aspects of the tissue distributions of a number of SLC transporters, which allows us to make the following general comments:

i. Even by inspection (although we also performed various tests for normality (Broadhurst and Kell, 2006), not shown), the distributions of transporters between different tissues or cell lines are very far from being normal, with instances of negligible expression in certain tissues for most transporters. The extreme here (and see below) is probably SLCO1B1 (Hagenbuch and Stieger, 2013), whose expression is virtually confined to the liver alone (a fact that has been exploited effectively for drug targeting purposes (Pfefferkorn, 2013; Pfefferkorn et al., 2012; Pfefferkorn et al., 2011));
ii. The tissue with the maximum overall expression of transporters (SLC and/or ABCs, total 11,331 TPM) is (probably unsurprisingly) the kidney (total 10,950, while that with the fewest is the pancreas (total 1,490 TPM) possibly explaining the difficulty of targeting drugs to it (Grixti et al., 2017));
iii. The SLCs with overall the greatest expression in total are SLC6A15 (a neutral amino acid transporter (Pramod et al., 2013), whose activity has been implicated in depression (Kohli et al., 2011)), and SLC25A3 (mitochondrial phosphate transporter (Palmieri, 2013)), while that least expressed *in toto* is SLC6A5 (a glycine transporter)
iv. The variation of expression is very considerable indeed. Almost every transporter ranges in its expression by over two orders of magnitude in different tissues, and several by more than three orders of magnitude (see also (Sreedharan et al., 2011; Winter et al., 2014)). We also show the minimum and maximum expression levels for each SLC, as well as the median and maximum.
v. The heatmap of expression levels (Eisen et al., 1998) shows a number of co-expression clusters, some of which we analyse below.
vi. Because the shape of the expression profiles varies so widely, we sought a suitable measure that would illustrate this. Measures such as inter-quartile range (normalised in any way) do not deal well with the extreme values, which are arguably of most biological interest. We decided that the Gini index (see Methods) was most suitable for our purposes.

**Figure 2.**
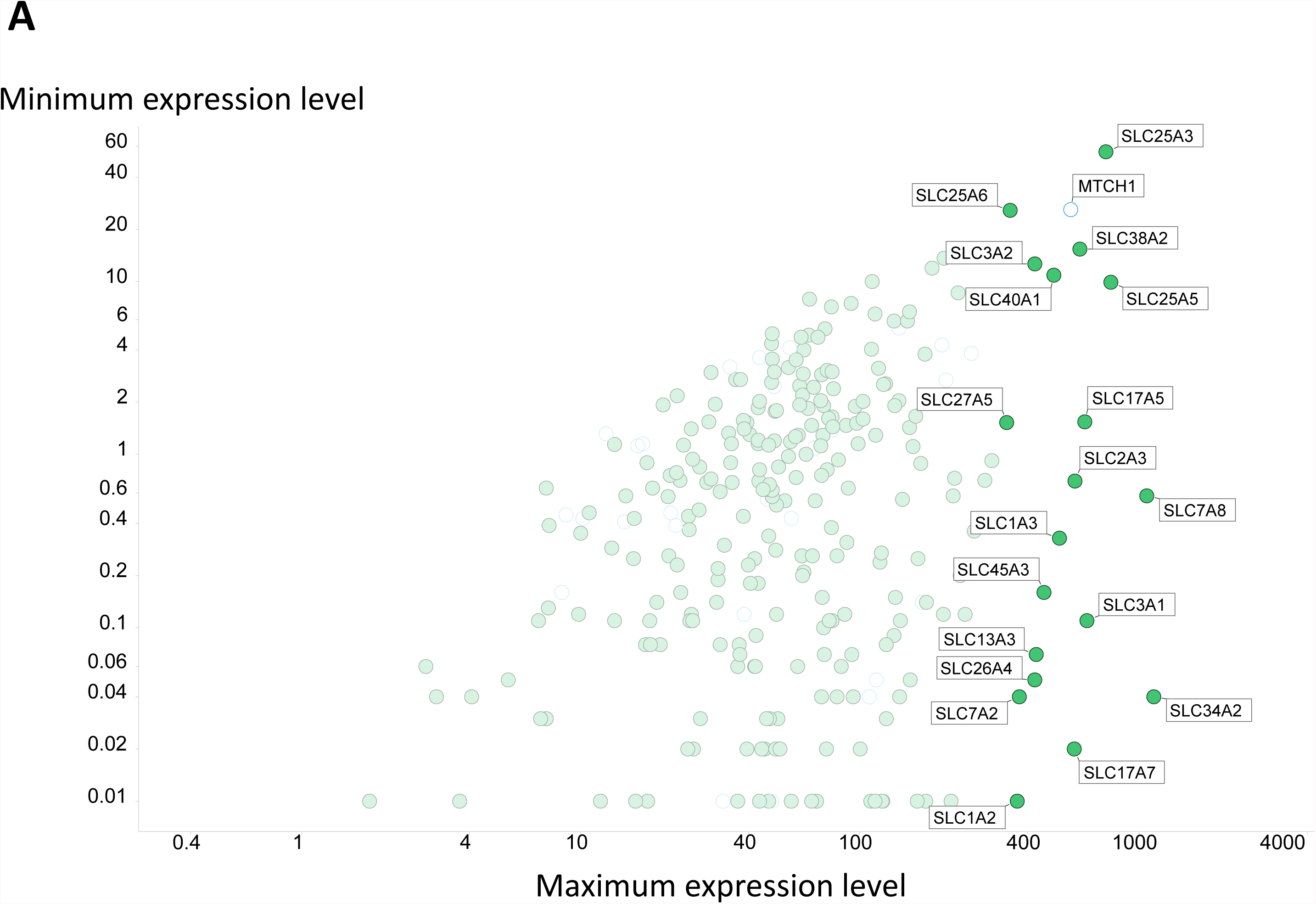
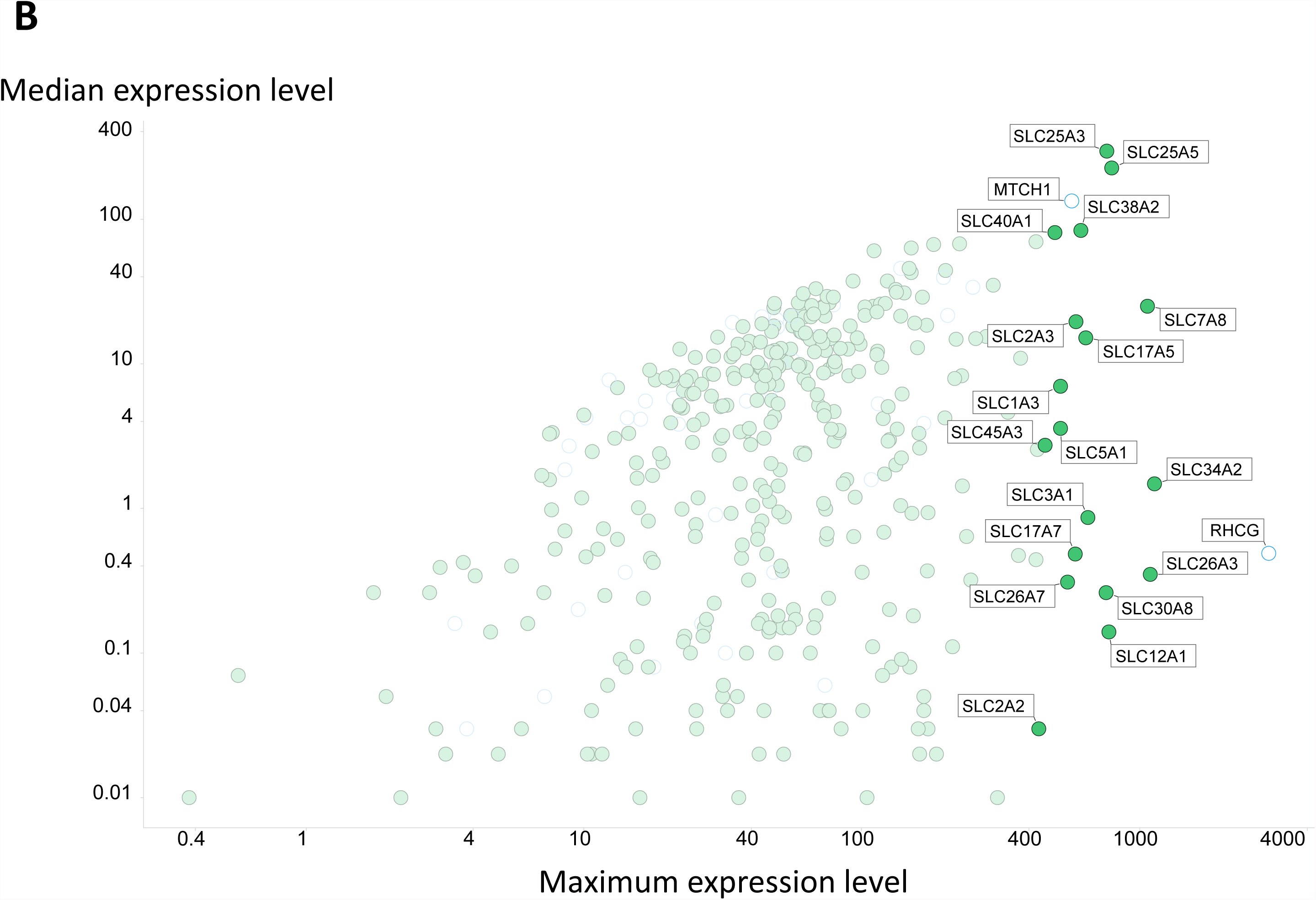
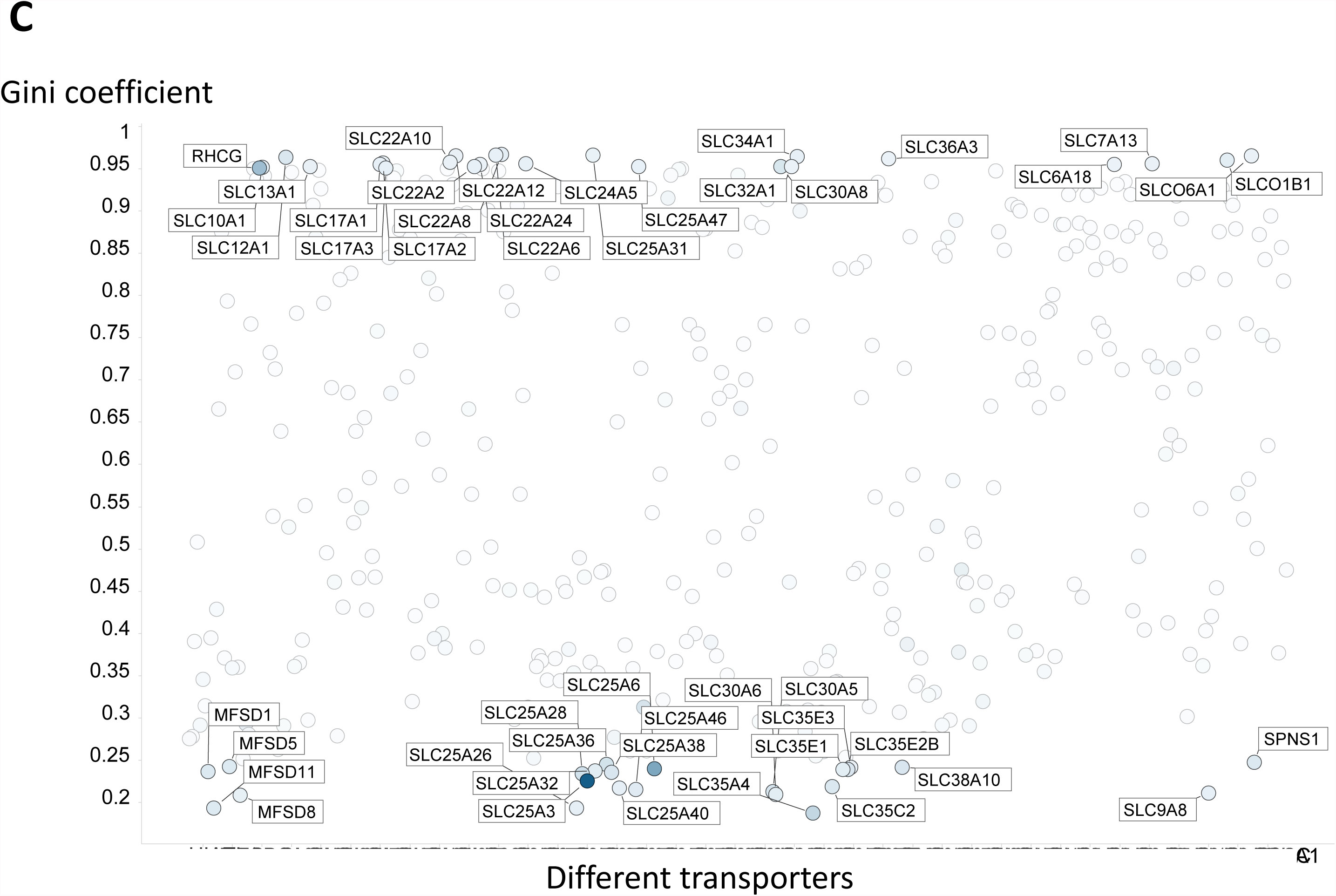
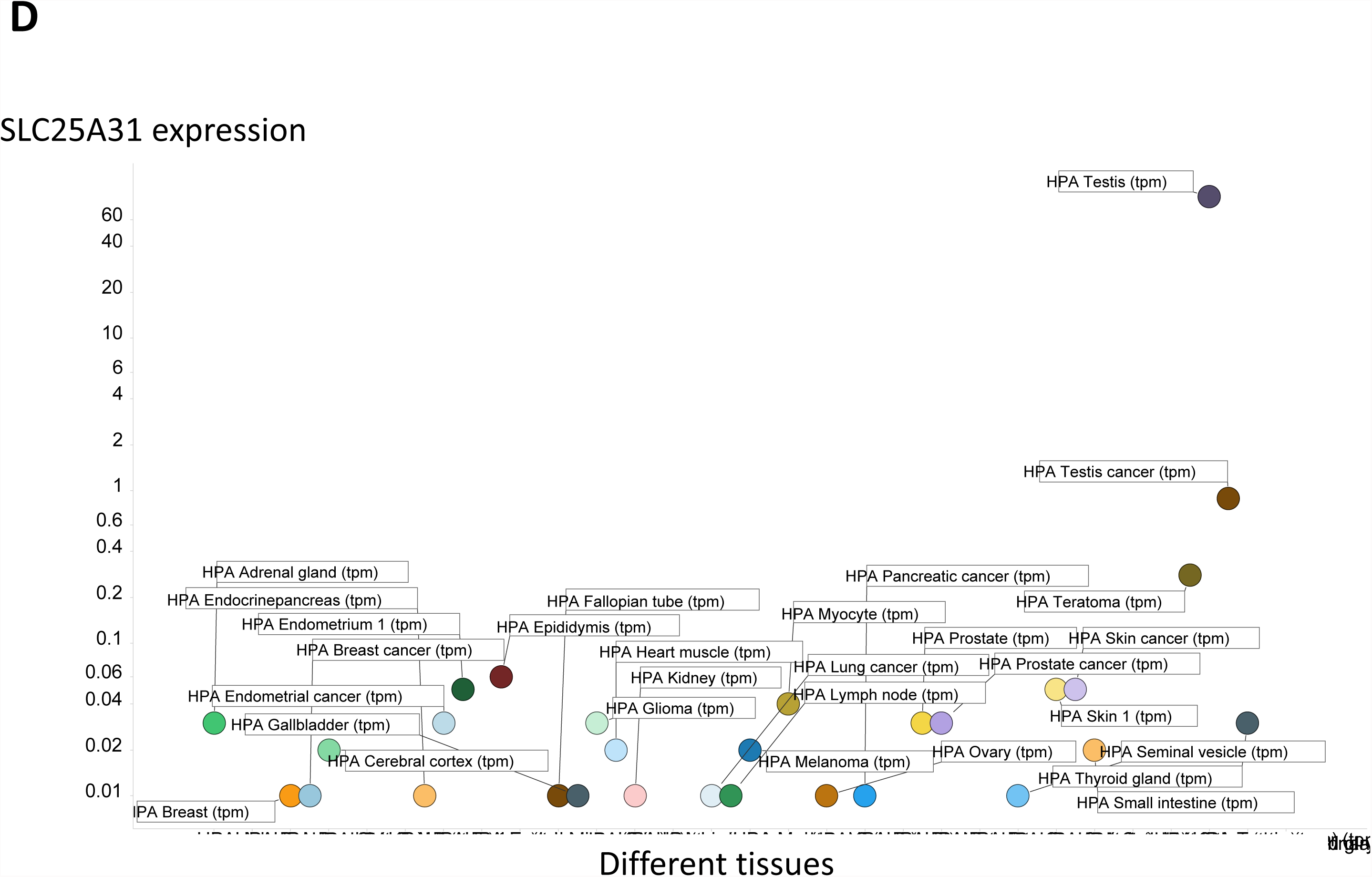
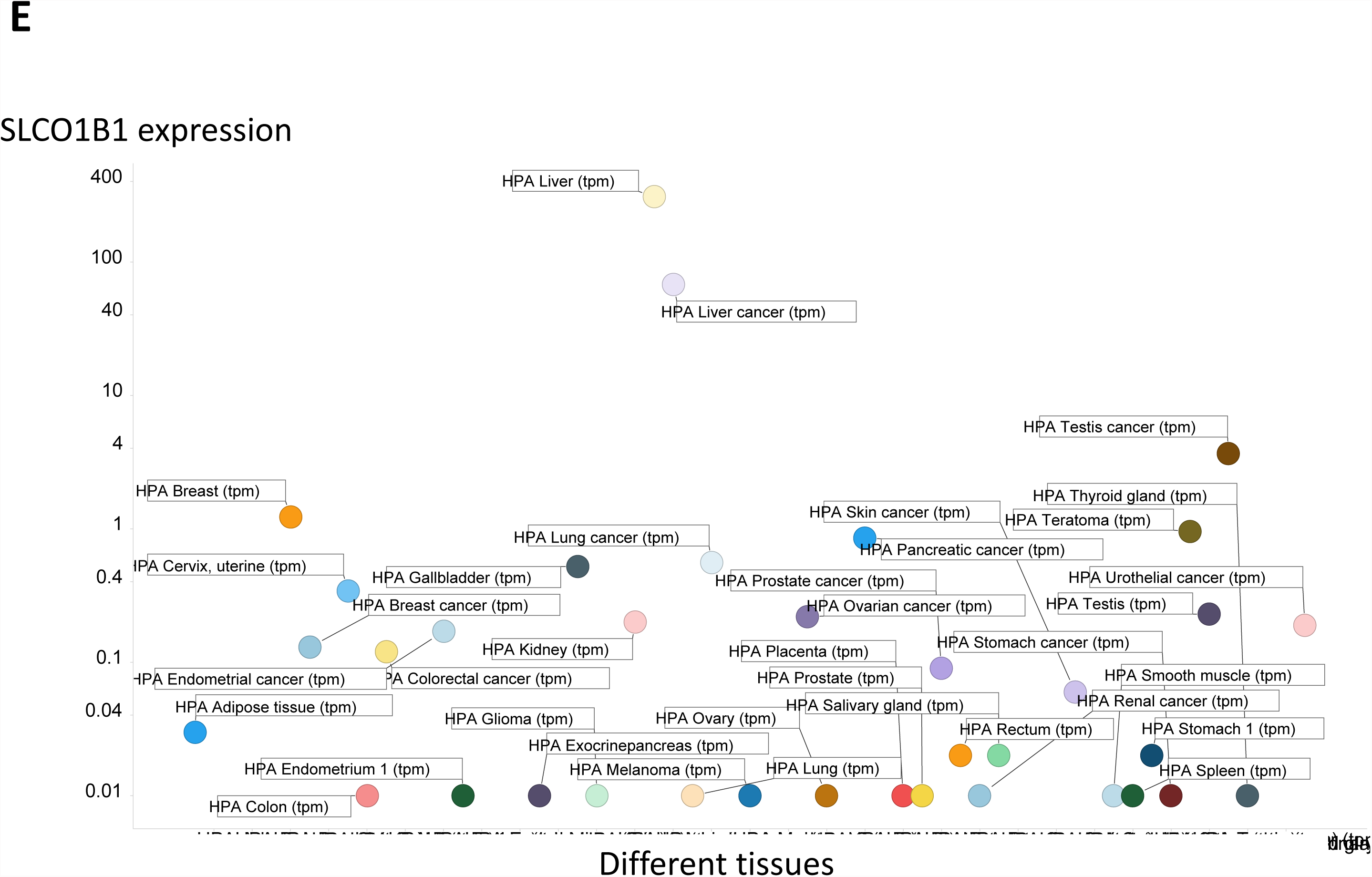
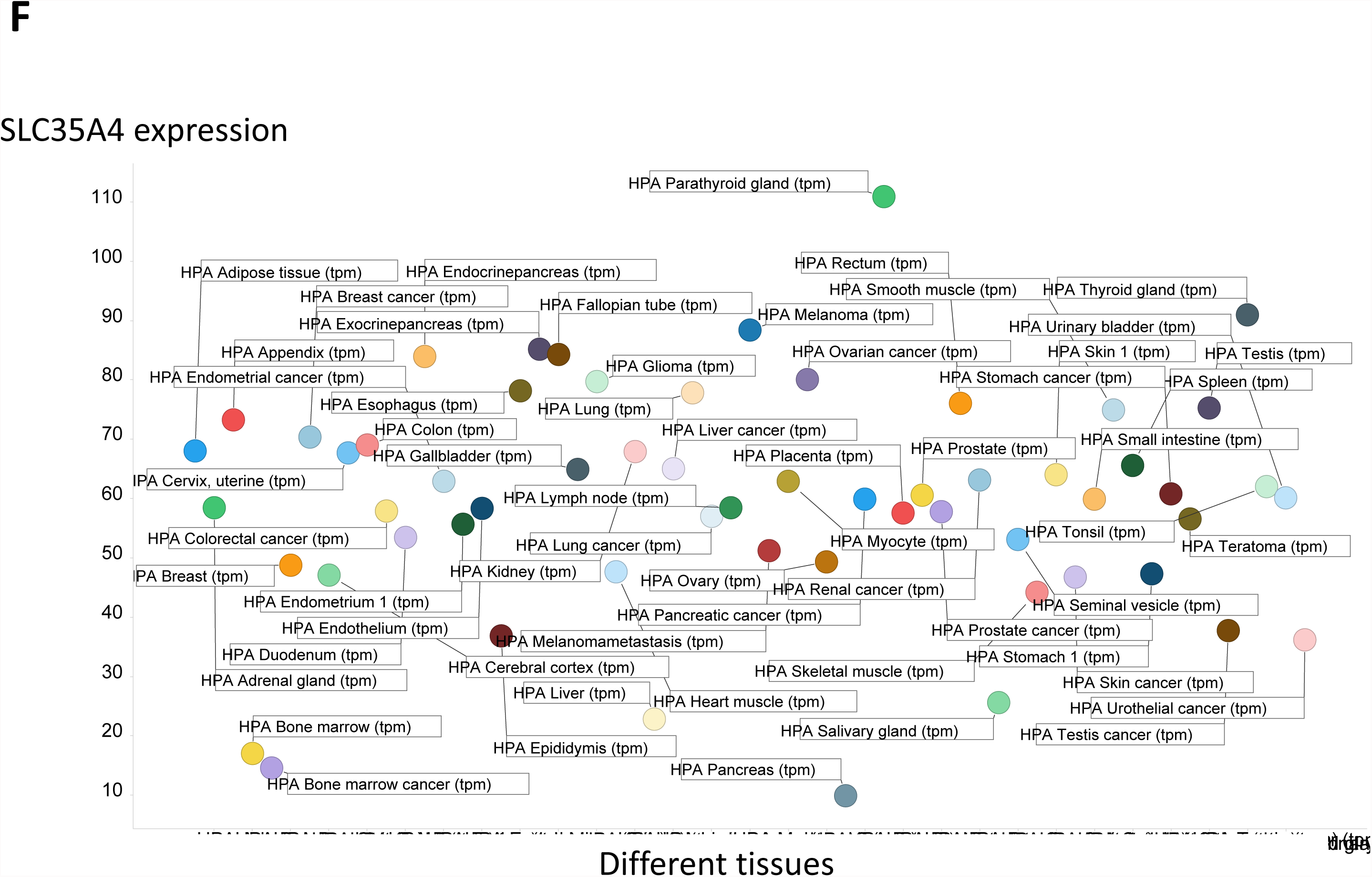
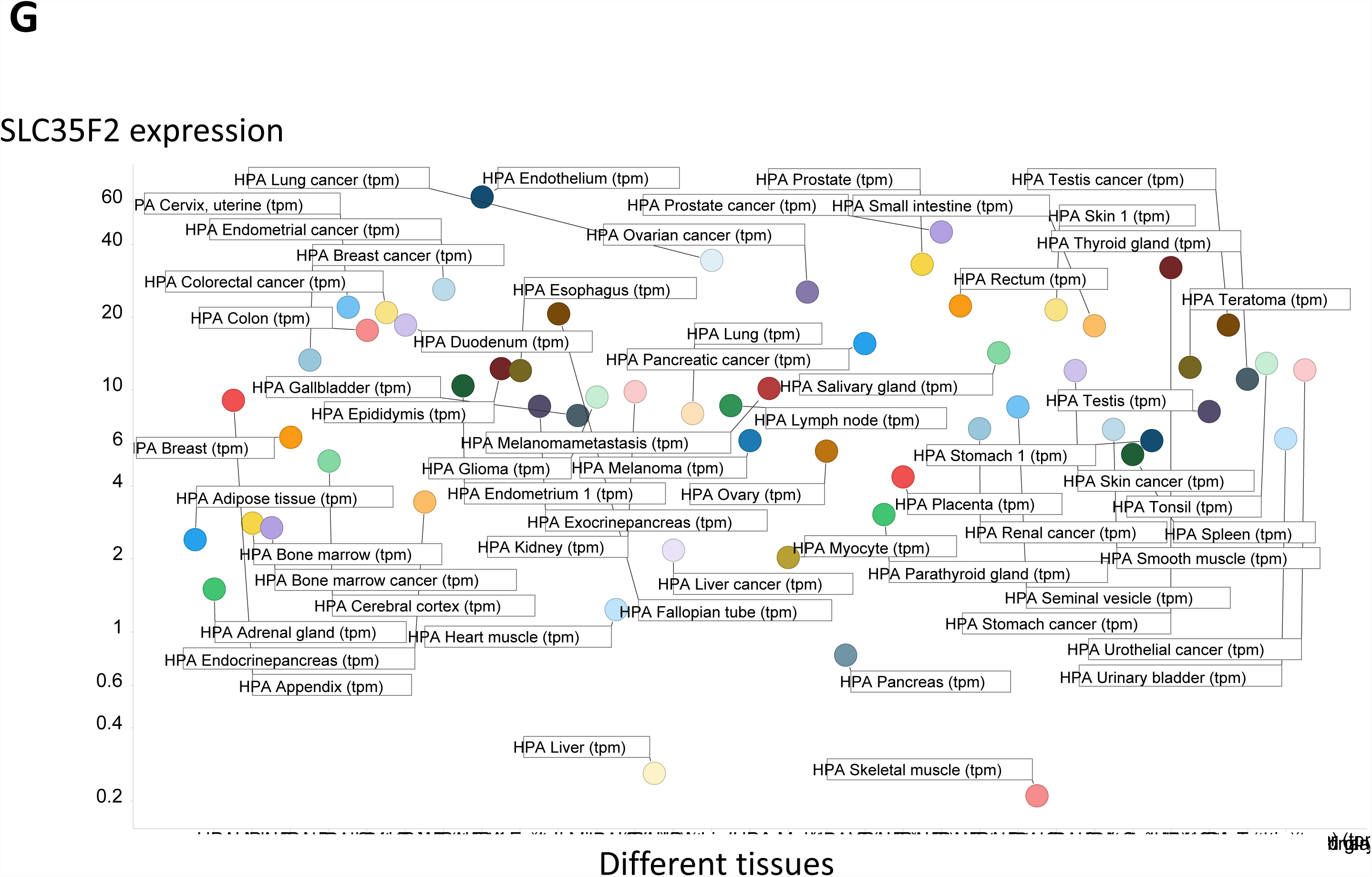
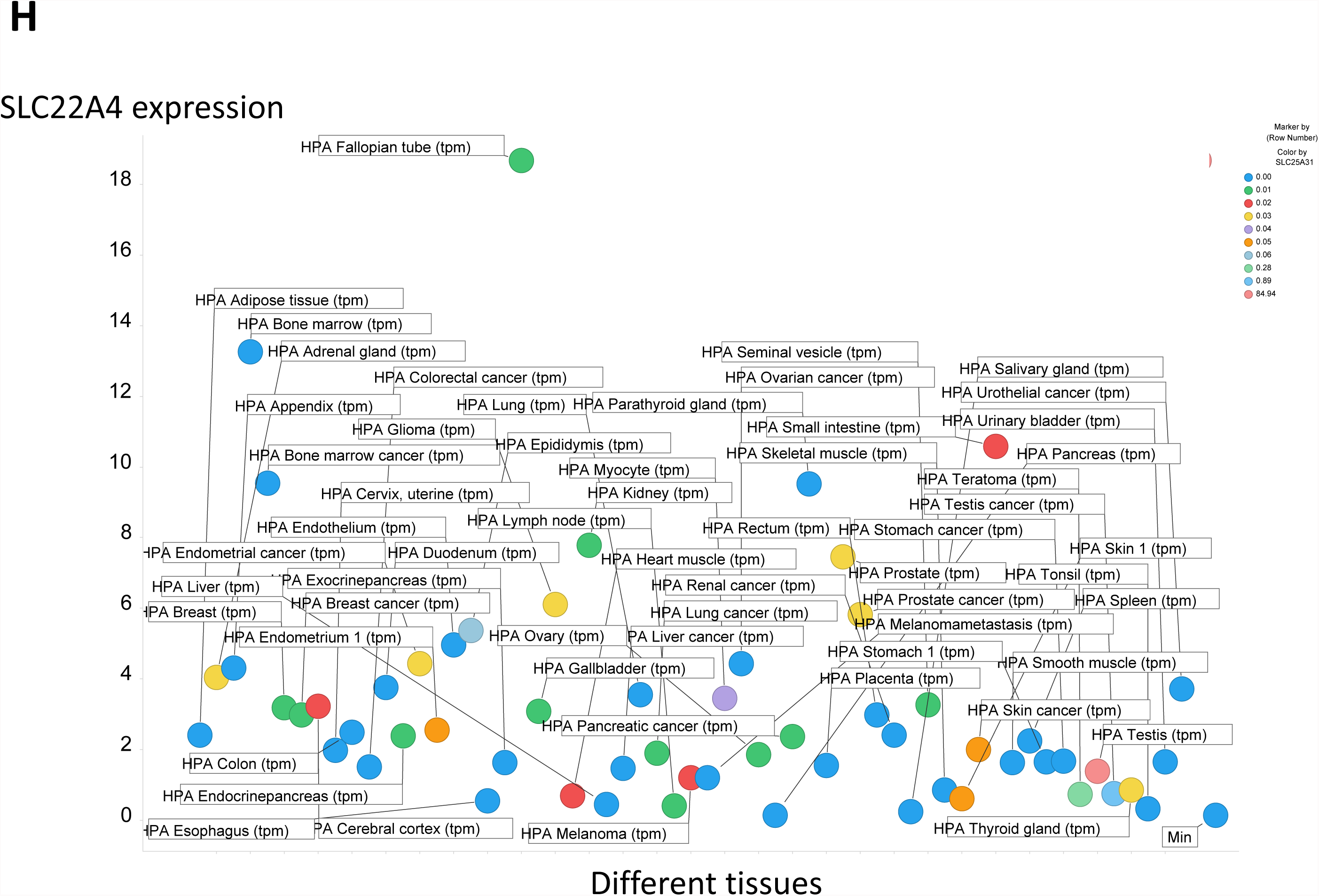
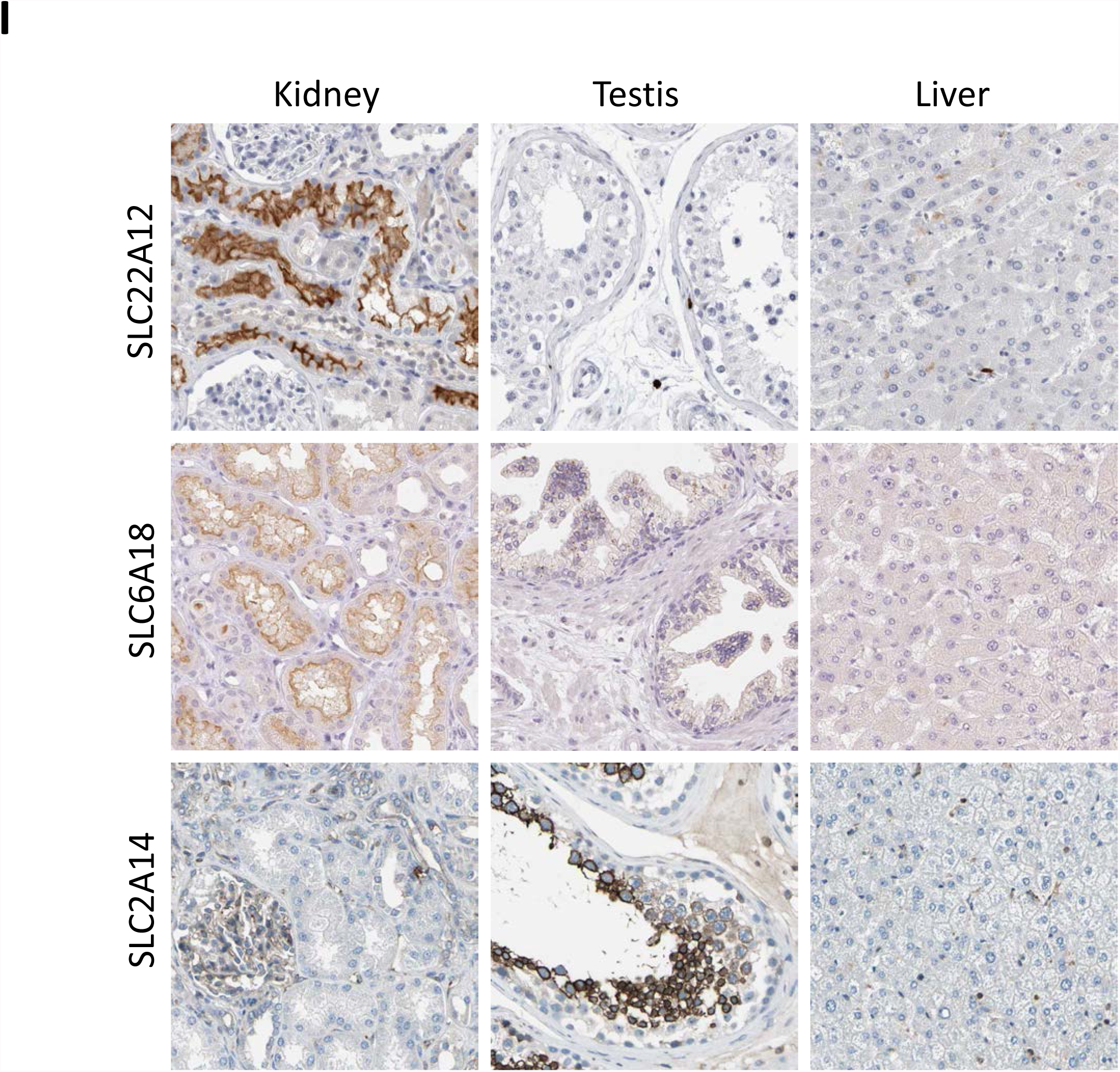
Expression profiling of various SLC transporters in 59 tissues. **A**. Minimum and maximum expression levels (ignoring those with undetectable expression) in the 59 tissues considered. **B**. Median and maximum expression levels (ignoring those with undetectable expression even at the median) in the 59 tissues considered. **C**. Gini coefficient for the expression of all SLCs in 59 tissues; those with Gini coefficients above 0.9 or below 0.25 are shown. **D**. SLC25A31 is almost exclusively expressed in the testes (the expression levels for others being 100x less). **E**. SLCO1B1 is almost exclusively expressed in the liver (with the expression level in other tissues being 100 times lower or less). **F**. The expression level of SLC35A4 is relatively homogeneous, with ¾ of all tissues within a factor two. **G**. The expression levels of SLC35F2 vary much more considerably, by a range of ~200 in these 59 tissue types. **H**. Expression profile of the transcripts for SLC22A4. **I**. Antibody-based expression of the SLC22A12, SLC6A18, and SLC2A14 transporters in kidney, testis and liver tissues. SLC22A12 and SLC6A18 are expressed in renal proximal tubules, whereas SLC2A14 is expressed in cells in seminiferous ducts. Image edge length is 320 μm.

Thus, Fig 2A shows the minimum and maximum expression levels (as TPM) for each transporter, with the top 20 (maximum expressions) labelled explicitly. Open circles are those not explicitly labelled as SLC family members. Interestingly, the mitochondrial transporters (Palmieri, 2013) SLC25A3 (for phosphate) and SLC25A5 (adenine nucleotide translocase (Clémençon et al., 2013)) are among the most highly expressed, as is the non-SLC MTCH1 which as its name implies is a MiTochondrial Carrier Homologue. The co-expression of SLC25A3 and SLC25A5 is entirely logical, as ATP synthesis and export requires the transport of equimolar amounts of its substrates. Many other SLC25 (mitochondrial transporter) family members are well represented as high expressers in at least one tissue. Note that expression levels below 0.01 TPM are not shown. Fig 2B shows similar data for the median versus the maximum expression in the different tissues. The median of the set of median expression levels for all the SLCs was 3.19. Thus, almost all transporters (404/409) have an expression level in at least one tissue that is above the median for each transporter between tissues; it is not at all the case that a transporter tends to be either highly expressed or weakly expressed. In other words, although many transporters are widely distributed, there is a considerable degree of specialisation (see also (Sreedharan et al., 2011)).

The Gini index for the variation in (or inequality of distribution of) transporters (Fig 2C) is fully consistent with this, with a significant number having an exceptionally high value (66 at 0.9 or above), not least SLC22 family members, often in the kidney (see below), and with only a few of the transporters (23/409) having a Gini coefficient below 0.25. One interpretation is that, mostly, individual transporters may be quite specialised; another is that different tissues require different amounts of specific substrates, though such large differences are not easily thereby explained in general. The <u>median</u> Gini coefficient for this overall class of SLCs and related transporters is 0.587 (higher than that of any country’s income distribution!). A number of those with the lowest GCs are again in the SLC25 (mitochondrial transporter) family; this is not unreasonable, since every cell is likely to have mitochondria, but some family members are clearly very specialised for particular mitochondria. Thus (Fig 2D) SLC25A31 (AAC4), a particular isoform of the adenine nucleotide translocase (Palmieri, 2013)), is essentially expressed only in the testes (Dolce et al., 2005) (Gini coefficient 0.965), a finding of unknown biological significance (Hamazaki et al., 2011). However, since its removal inhibits spermatogenesis (Brower et al., 2007) and thus causes infertility (Brower et al., 2009), it is potentially a target for the development of male contraceptives.

As mentioned, SLCO1B1 (a major transporter of so-called statins) is confined essentially only to expression in the liver (Fig 2E), and its Gini coefficient is ~0.96. At the other end of the Gini spectrum (Gini coefficient = 0.188), transporters such as SLC35A4 are almost universally expressed (Fig 2F). However, this is not true of all SLC35 family members, since SLC35F2 enjoys a very wide distribution of expression levels in both tissues (Fig 2G) and cell lines (Winter et al., 2014). We also have an interest in the ergothioneine transporter (SLC22A4, previously known as OCTN1) (Gründemann, 2012; Gründemann et al., 2005); Fig 2H shows its expression profile distribution in the tissues considered; its Gini coefficient is 0.502. A much more restricted analysis of its expression profile was given by Gründemann and colleagues (Gründemann et al., 2005), and it did not include the Fallopian tube (the tissue with the greatest expression level here). Finally, we illustrate (Fig 2I) the expression of SLC22A12 (URAT1, a urate transporter) (Anzai et al., 2012; Koepsell, 2013; Pelis and Wright, 2014) in the kidney, virtually the only tissue in which it shows expression (Gini index = 0.978).

One hypothesis might be that nutrient transporters (Palm and Thompson, 2017), e.g. those for glucose (part of the major facilitator superfamily (Reddy et al., 2012)), or for amino acids (Bröer and Palacín, 2011; Taylor, 2014) (part of the amino acid/polyamine/organocation (APC) superfamily (Höglund et al., 2011; Jack et al., 2000)), might be more universally expressed, but this does not seem to hold up. Thus, SLC6A18, a neutral amino transporter, has the 15^th^ highest Gini coefficient (0.955), and its expression is essentially confined to the kidney proximal tubule. Similarly, SLC2A14, a glucose transporter (Mueckler and Thorens, 2013), has a GC of 0.853 and is again largely confined to the testes (Fig 2I). Mueckler and colleagues comment (Mueckler and Thorens, 2013), however, that its physiological substrate is unknown, despite it having 95% sequence identity to the *SLC2A3* gene. An overall theme is certainly that we do often lack knowledge of the ‘natural’ substrates of even well-studied drug transporters (e.g. (Gründemann et al., 2005; O’Hagan and Kell, 2017c; Perland and Fredriksson, 2017)).

### Correlations and heat maps

Some unexpected correlations arise, e.g. that between the expression of SLC39A5 (ZIP5, a Zn^++^ transporter (Jeong and Eide, 2013)) and SLC17A4 (supposedly a sodium/phosphate transporter in the vesicular glutamate transport family, of unknown function (Reimer, 2013); r^2^ = 0.86) (Fig 3A). Such findings raise many questions but provide few present answers. However, they do provide useful starting points for the testing of biological hypotheses.

**Figure 3.**
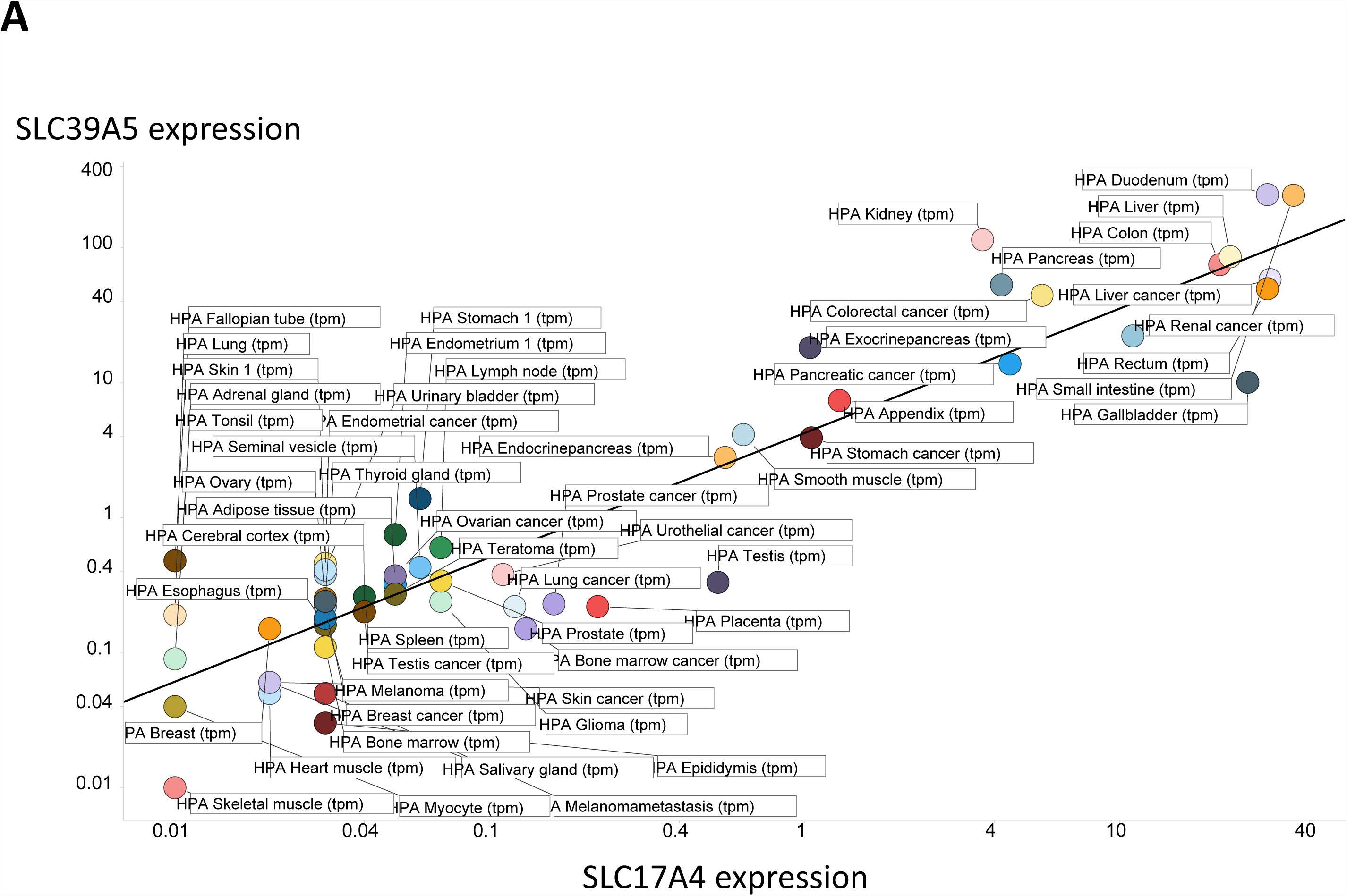
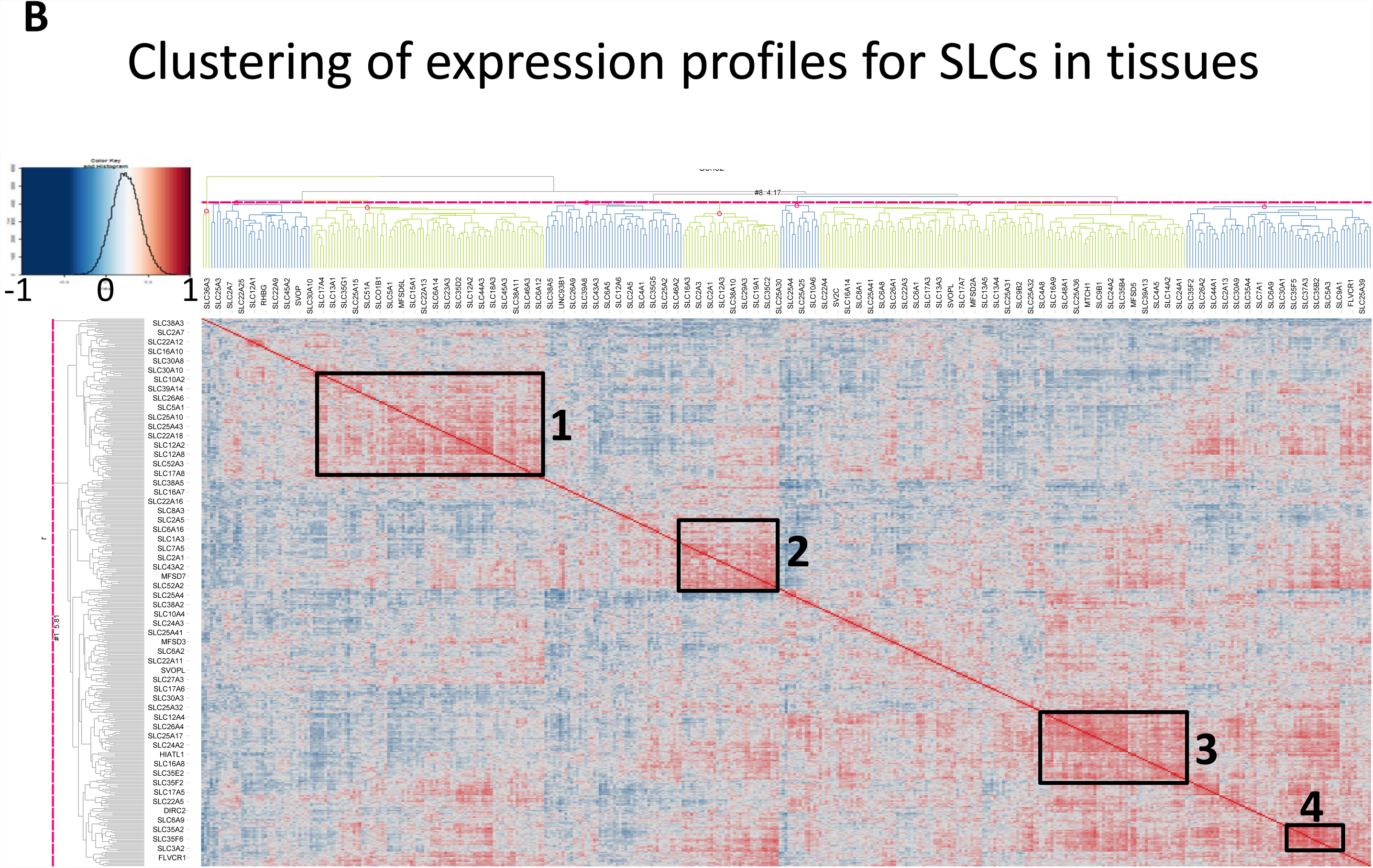
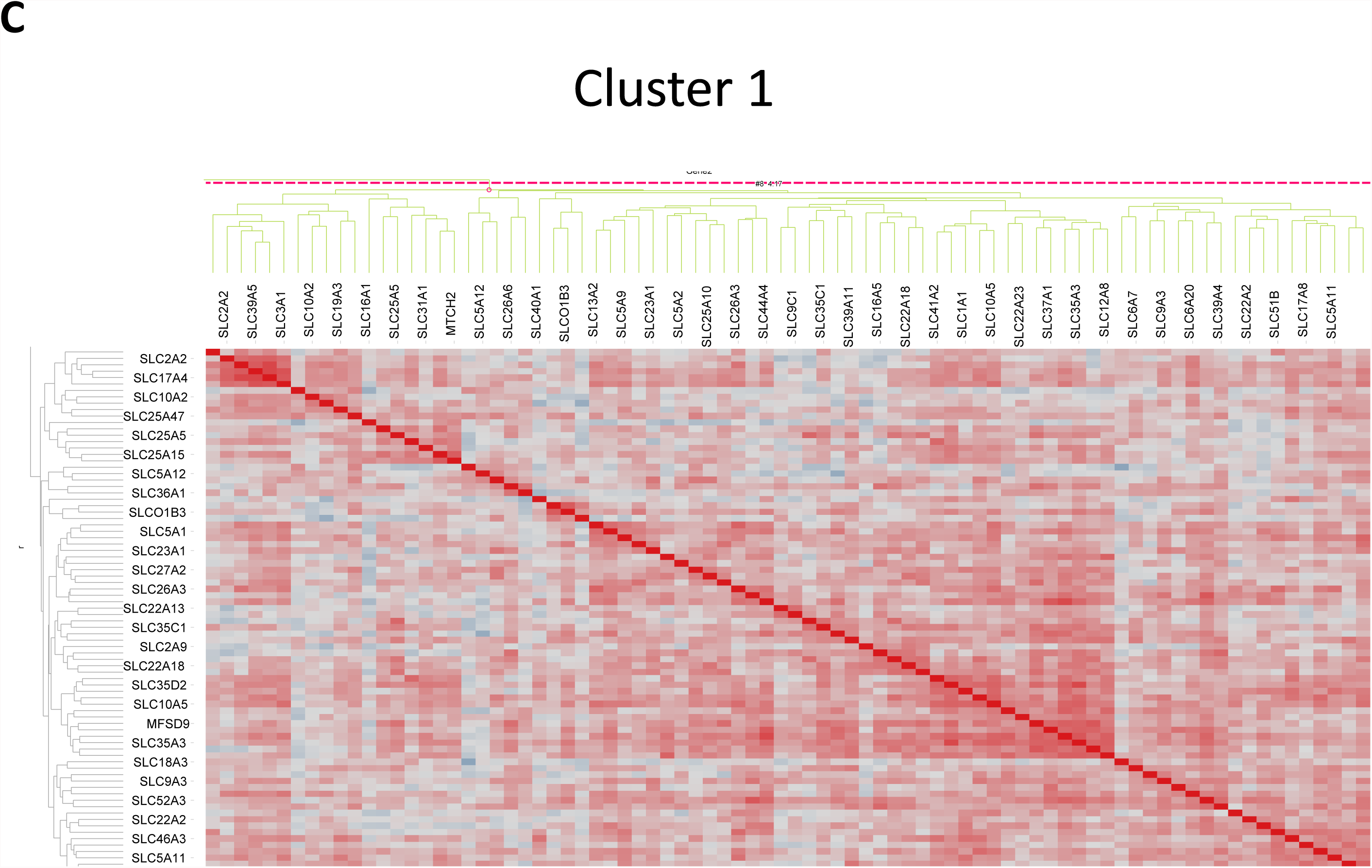
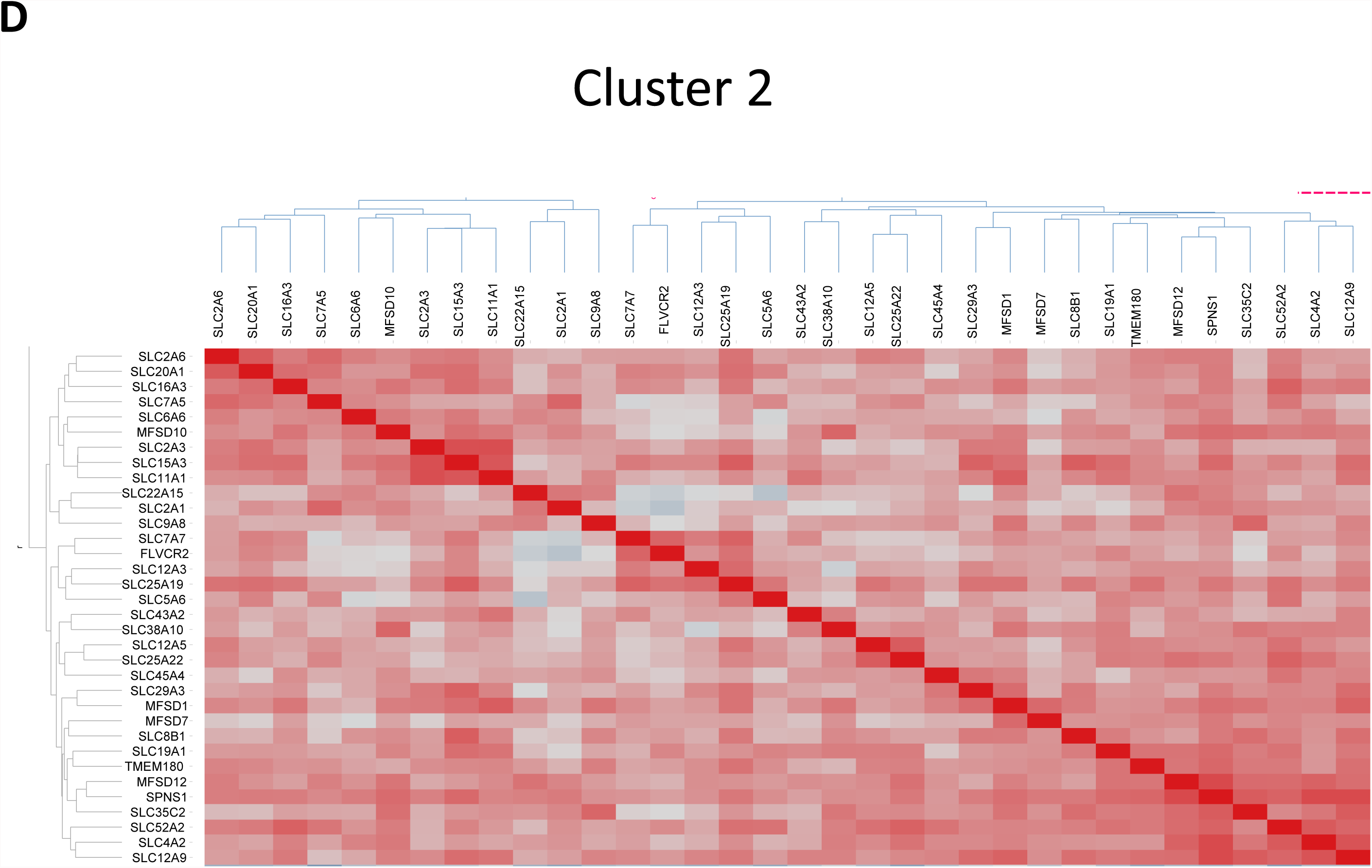
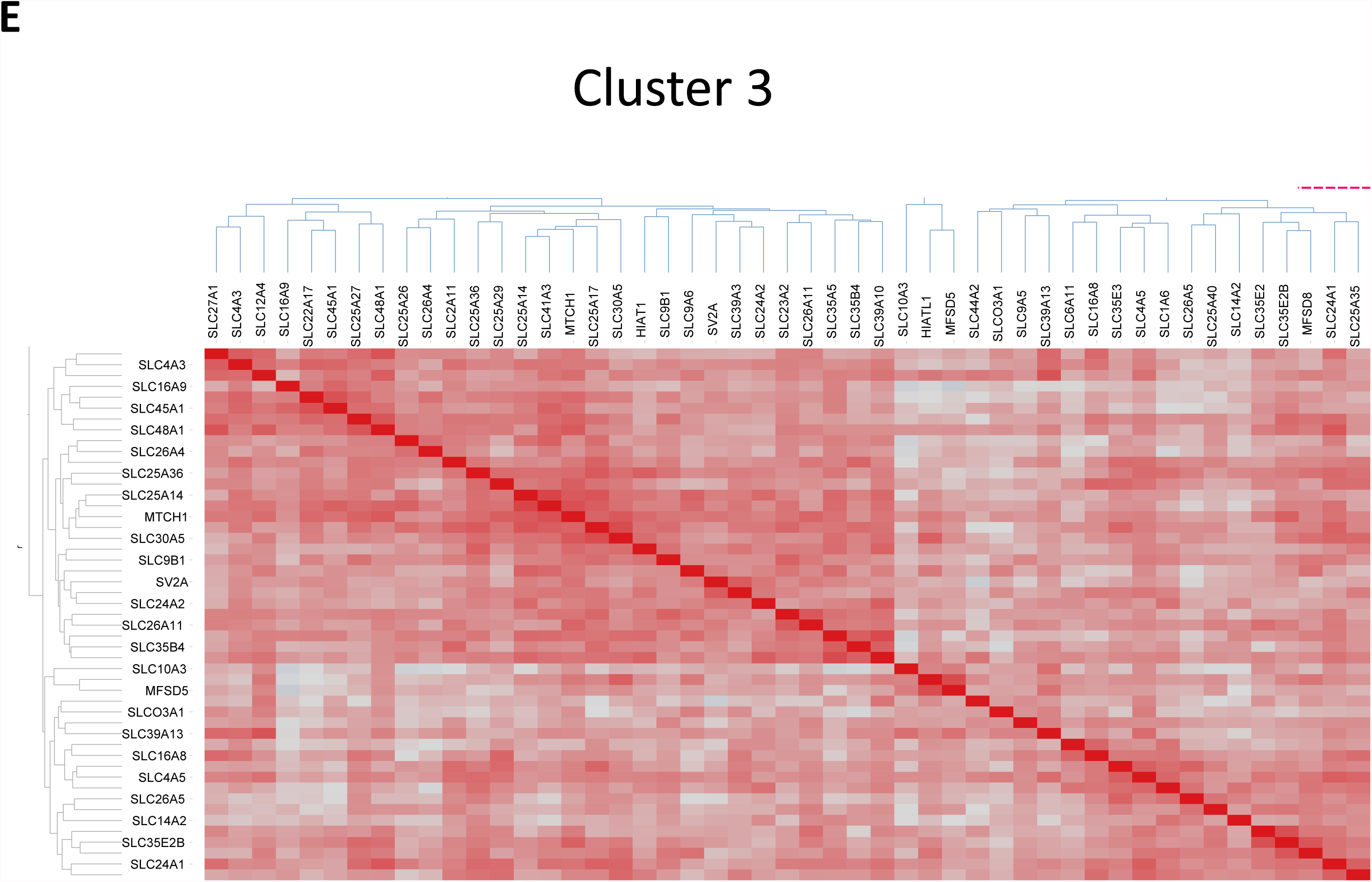
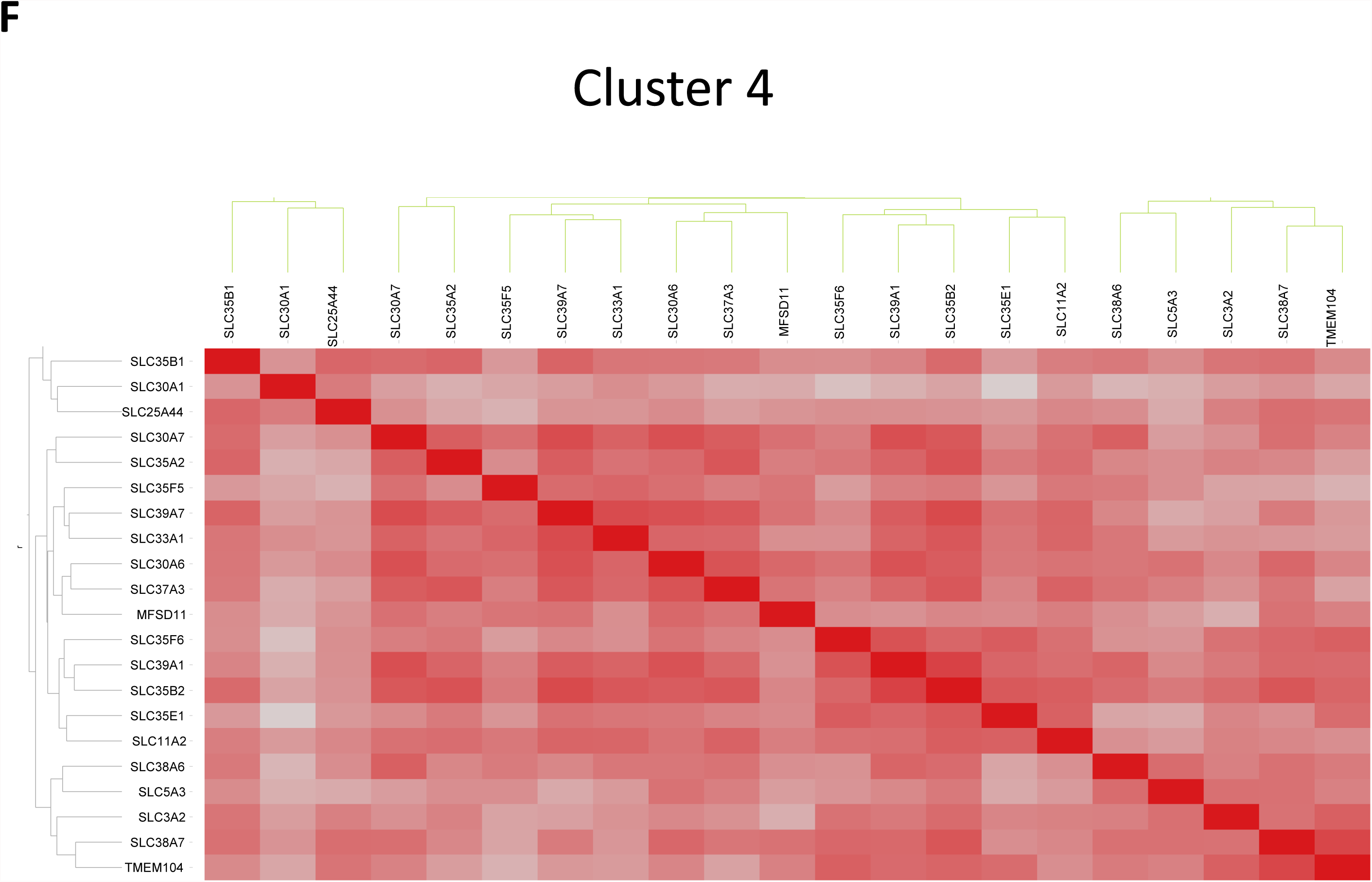
Clustering of (co-)expression profiles of SLC transporters. **A**. Significant correlation (in log-log space) between the expression profiles of SLC39A5 and SLC17A4 (r^2^ = 0.86). **B.** Overall heat map, with four major clusters highlighted. **C** to **F.** Zoomed-in versions of the four clusters.

In a similar vein, co-clustered heat maps of expression levels (Eisen et al., 1998) provide a very convenient visual summary of large amounts of data. Thus, Fig 3B shows the full heat map for SLC expression in tissues. We have marked four major clusters (zoomed in in Figs 3C-3F). With the exception of a slight preponderance of families SLC 25 and 35 in cluster 3 (Fig 3E) and of SLC35 in cluster 4 (Fig 3F), there was no obvious clustering at the level of families. This gives weight to the idea that SLC transporters have mainly exhibited <u>divergent</u> evolution (Höglund et al., 2011), in which similar sequences and structures acquire, adopt and are selected for different functions.

### SLCs in cell lines

A similar analysis was undertaken in a series of cell lines. Thus, as in Fig 1, we plot the minimum non-zero vs maximum expression levels (Fig 4A). The trends are broadly similar, with some of the most highly expressed transporters again being SLC25A3, SLC25A5, MTCH1, and SLC3A2, although there are also differences. The overall spread seems broadly similar to those of tissues, with a preponderance of transporters having minima in the decade 1-10 TPM and maxima in the decade 20-200 TPM. Overall, this might be thought of as giving confidence in the idea that cell lines were a reasonable representation of the behaviour of tissues, albeit their growth environment may have been significantly different. In a similar vein (Fig 4B), the number of SLCs with a Gini coefficient over 0.9 is 70, while those with GCs below 0.25 is 35. These numbers are also close to those for tissues, and the behaviour of the different <u>types</u> of transporters is again similar, with mitochondrial transporters tending to have a low GC. The median GC for SLCs in cell lines (0.595) is very close to that for tissues (0.587). This again implies that the extent of differentiation of transporter expression profiles between the tissues and the cell lines examined is broadly similar, as expected for established cell lines (Masters, 2000), albeit the range is greater in the former. This may be a result of the mixture of cell types in the tissues and that some (or even many) transporters likely exhibit a cell type-specific expression pattern like SLC22A12, SLC6A18 and SLC2A14 (Fig 2I). Finally (Fig 4C) we show the extensive variation in expression profiles of SLC22A4 (the ergothioneine transporter) in the different cell lines.

**Figure 4.**
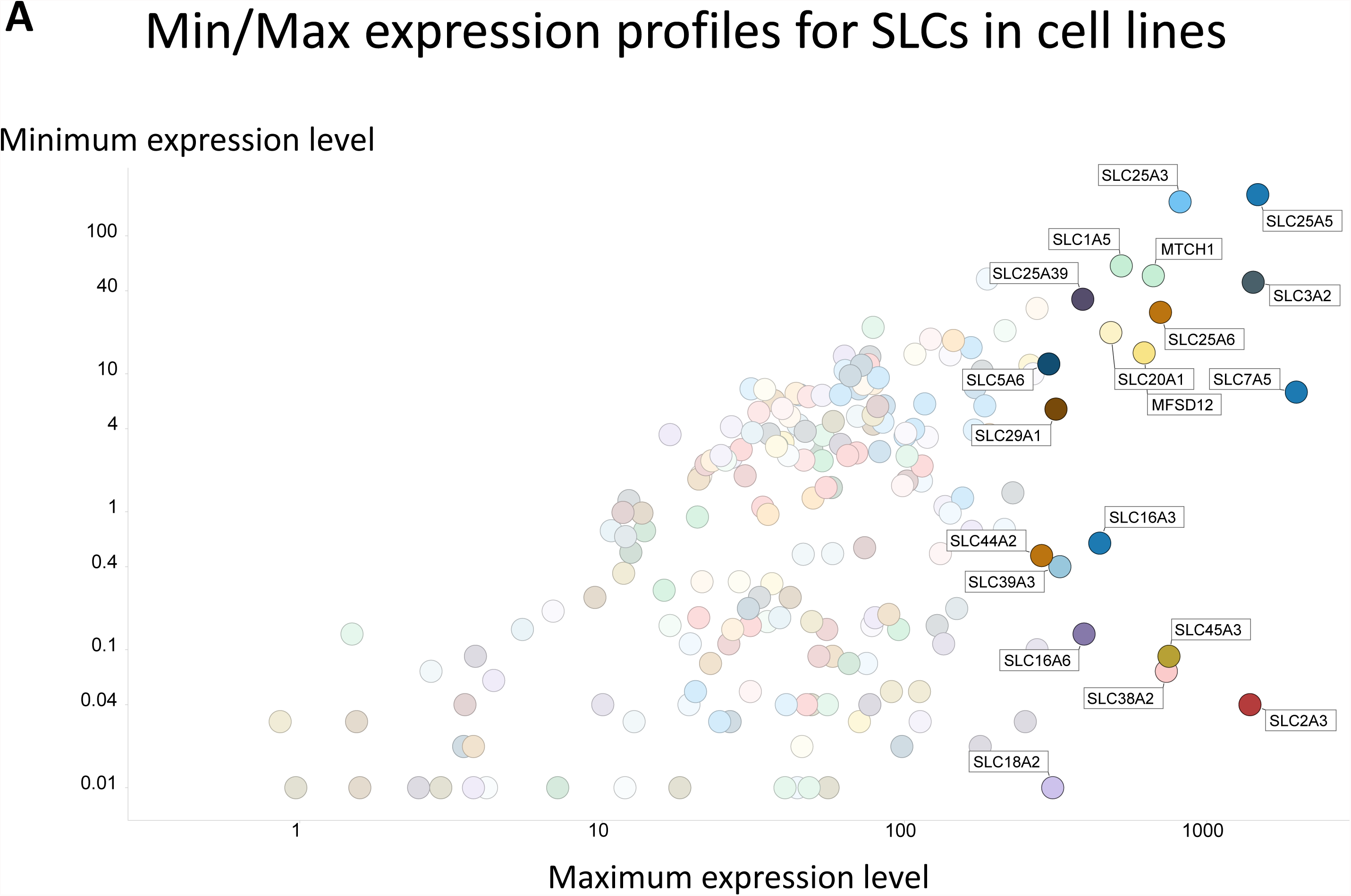
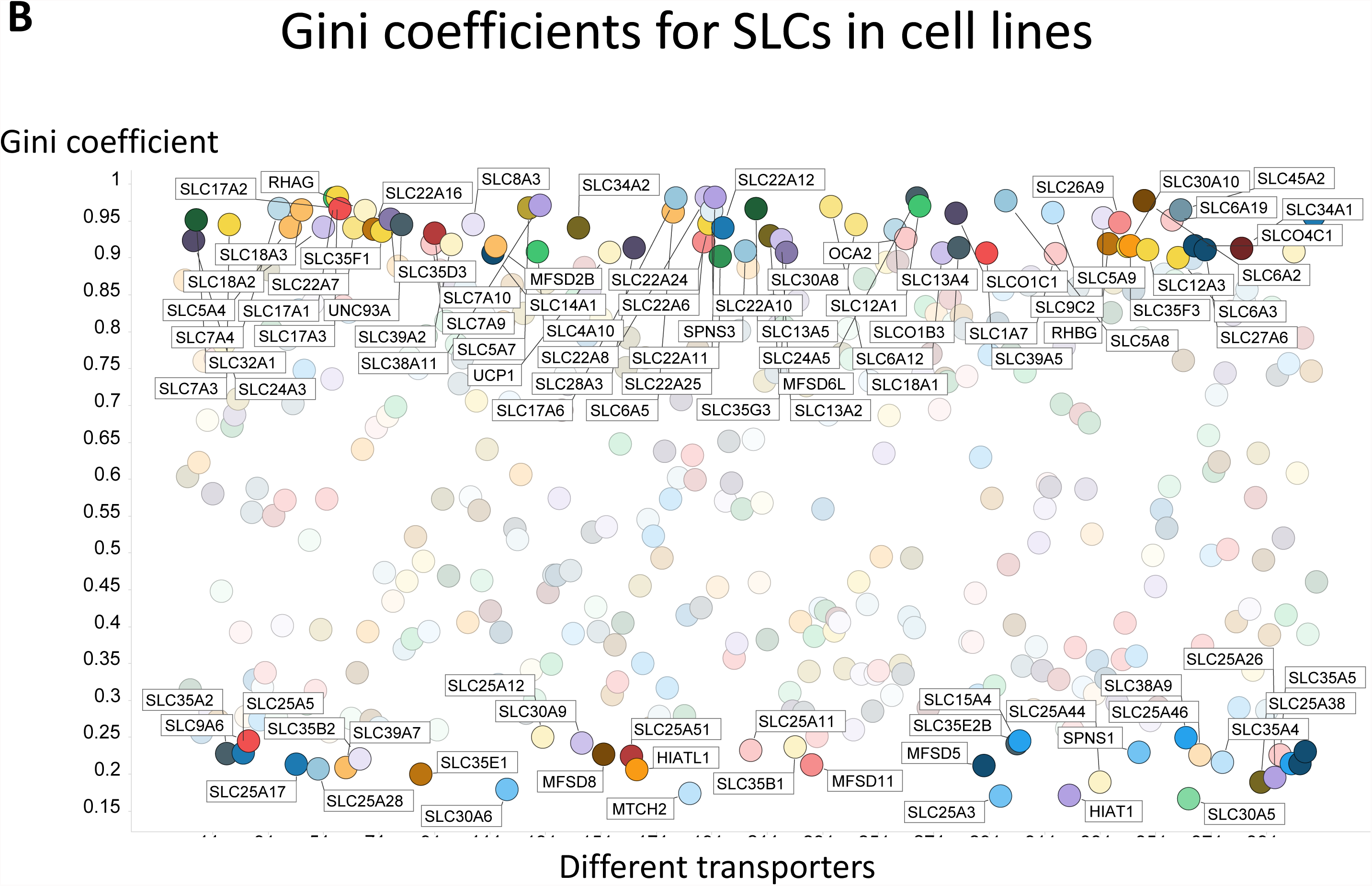
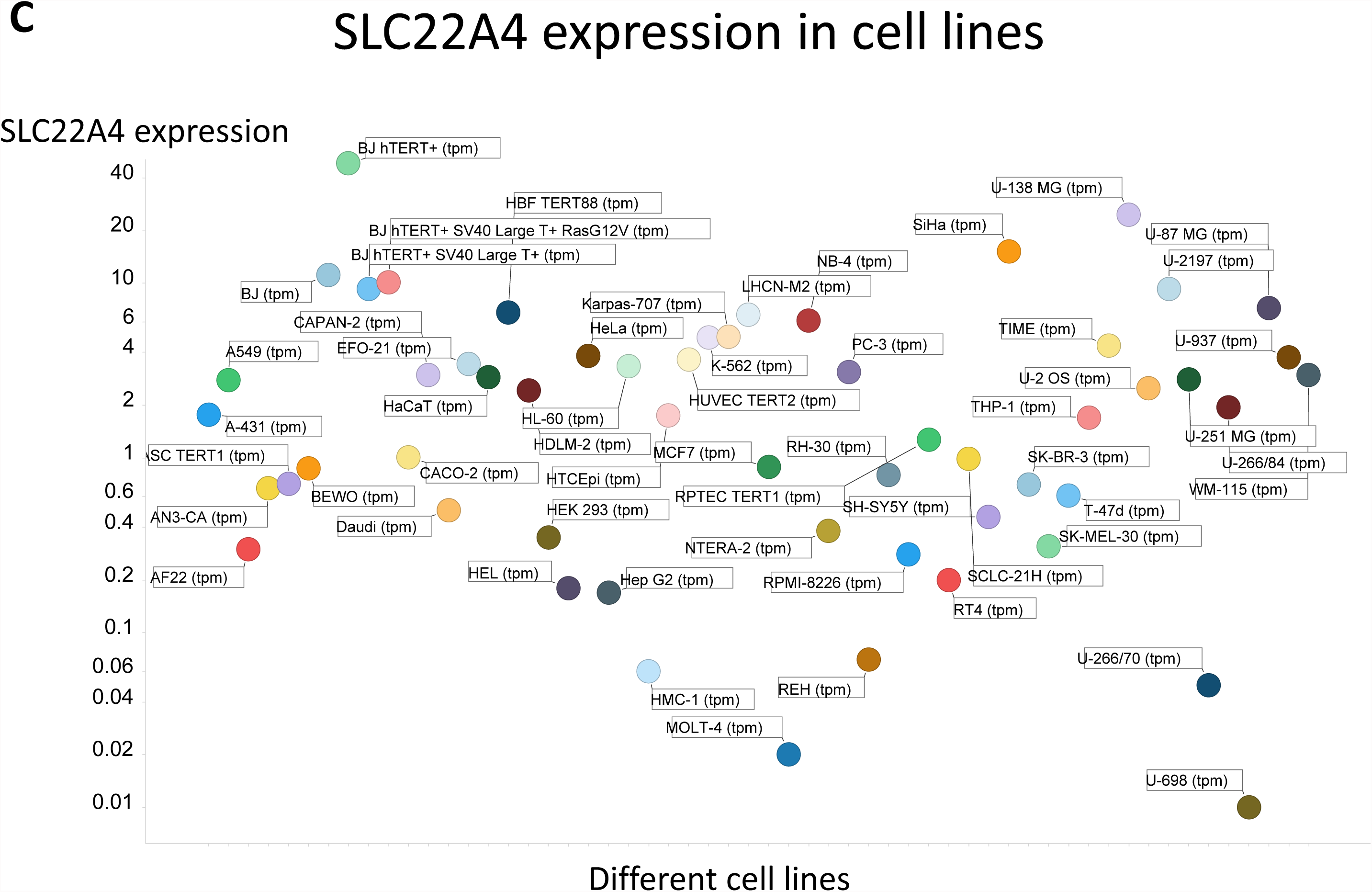
Expression profiling of various transporters in 56 cell lines. **A**. Minimum and maximum expression levels (ignoring those with undetectable expression) in the 56 cell lines considered. **B**. Median and maximum expression levels (ignoring those with undetectable expression even at the median) in the 56 cell lines considered. **C.** SLC22A4 expression levels (in TPM) in different cell lines.

### ABC transporters in tissues

Figure 5A shows the minimum and maximum expression levels for all 48 ABCs, many of which lack detectable expression in at least one tissue type. Again, the <u>ranges</u> of expression are considerable, but the expression levels of the ABCs tend to be slightly lower than are those of the SLCs in tissues. The total numbers are small, but no family (encoded in colour in Fig 5A) except possibly F seems especially highly expressed. The overall most highly expressed ABC transporter is ABCC4. The Gini coefficients (Fig 5B) vary more than do those of the SLCs, and have a median value of 0.496. 5/48 GCs are greater than 0.9, while four are below 0.25. Several ABCs exhibit very high Gini coefficients, that (0.939) of ABCG5 being the largest; it is mainly expressed in the duodenum and the liver. Those of the F family, however, while highly expressed, also have a low Gini coefficient, indicating that they tend to be among the more highly expressed in most tissues. Indeed, consistent with this, they are probably not in fact transporters (Dean and Annilo, 2005; Nishimura et al., 2007; Tyzack et al., 2000).

**Figure 5.**
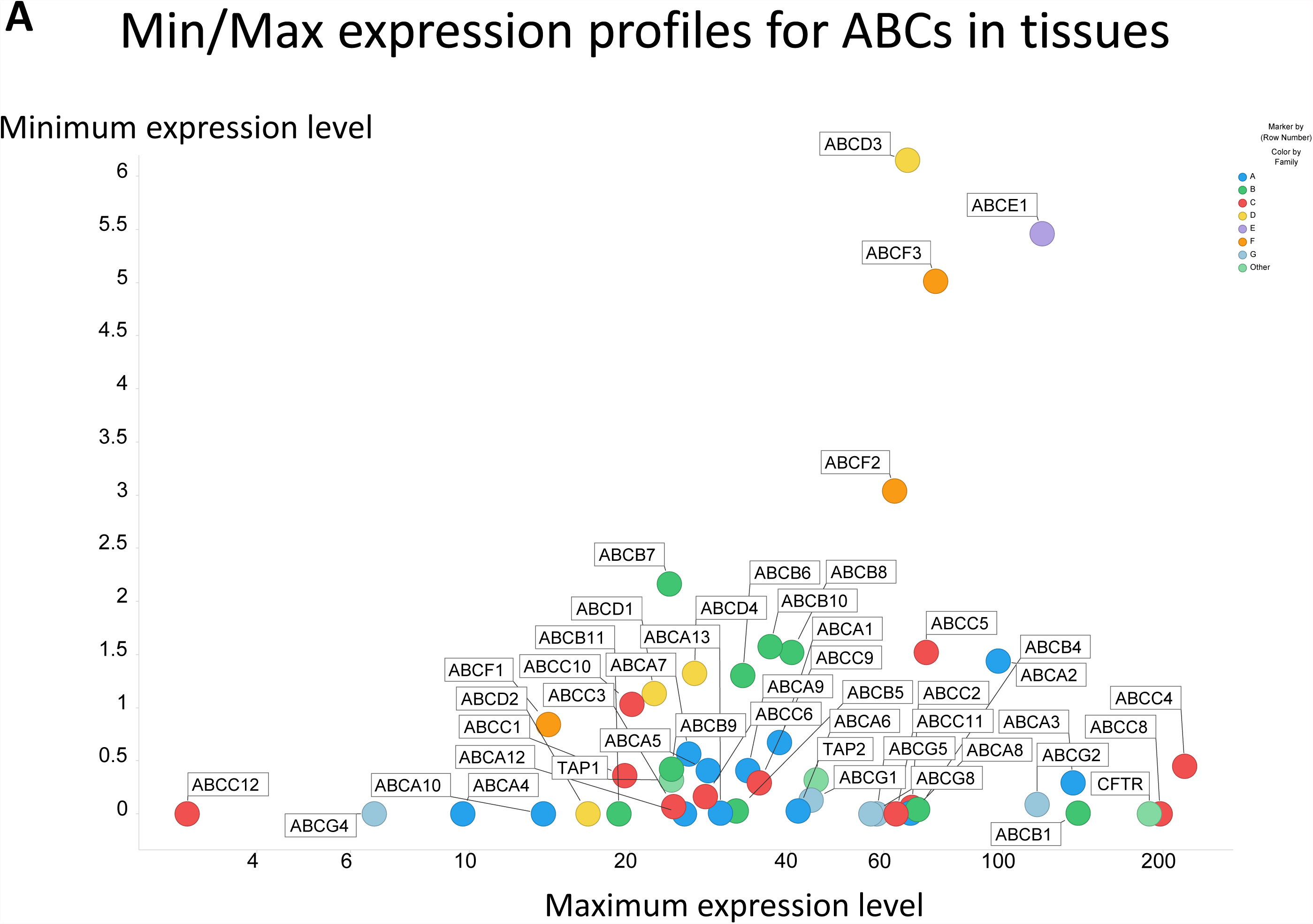
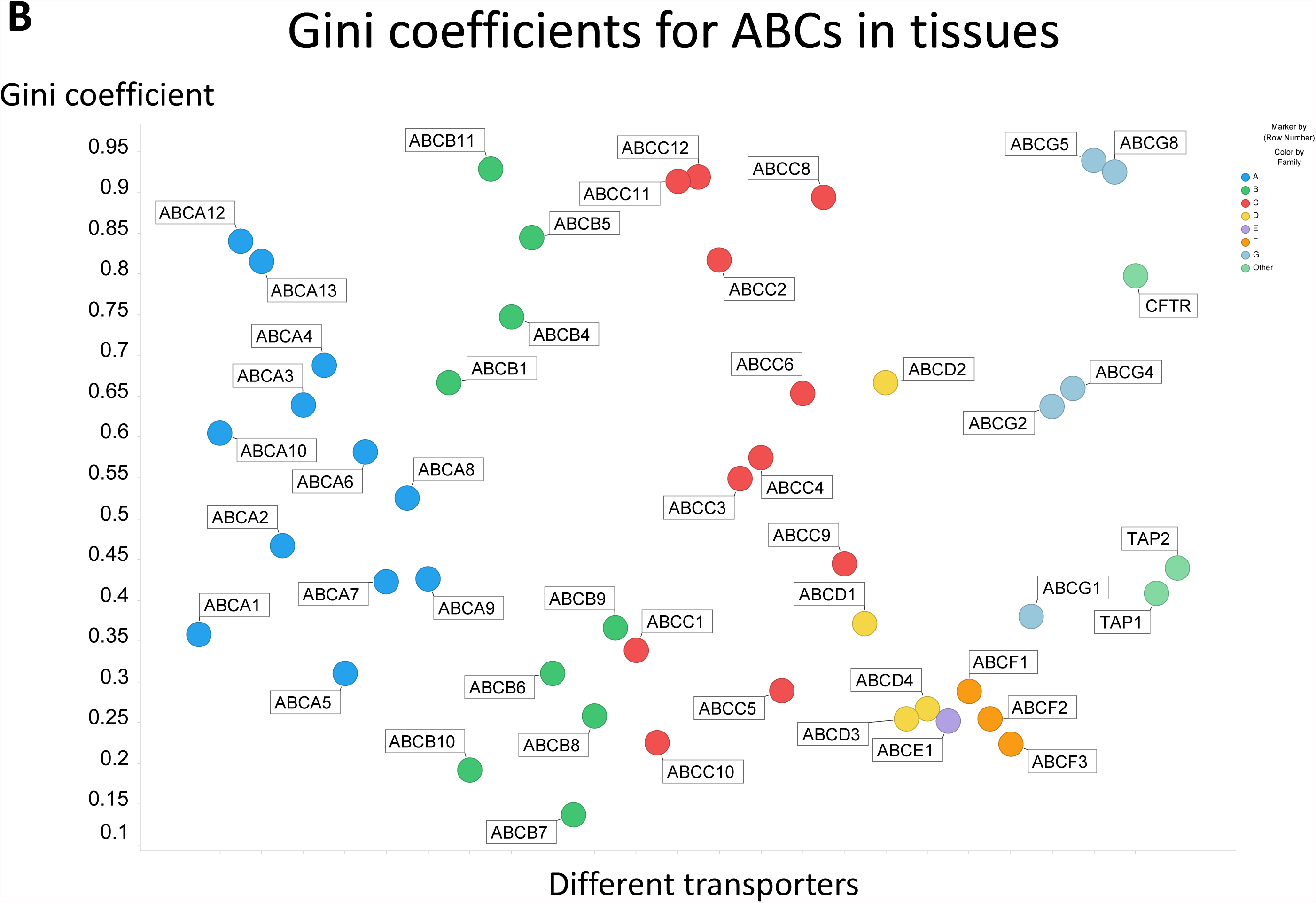
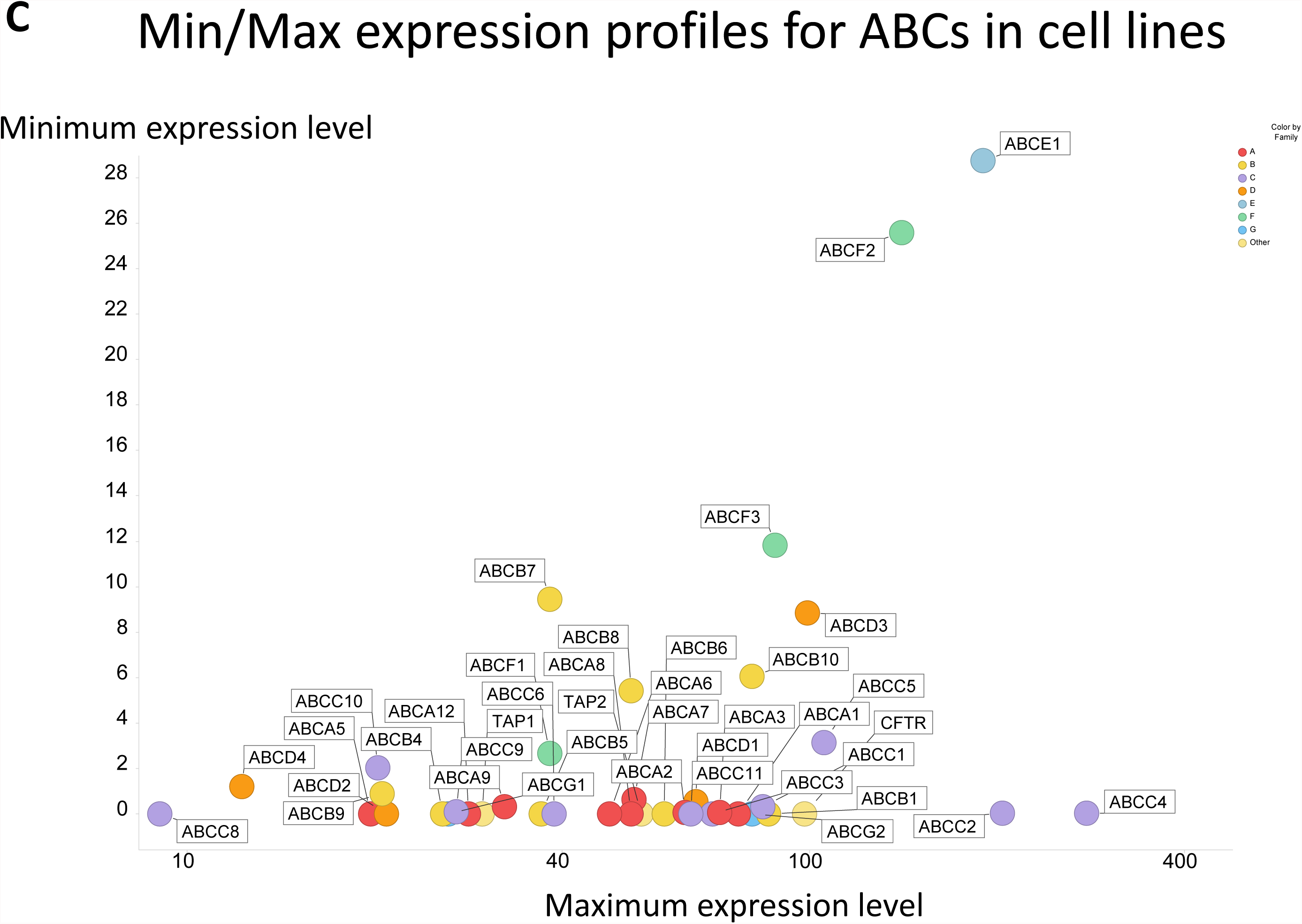
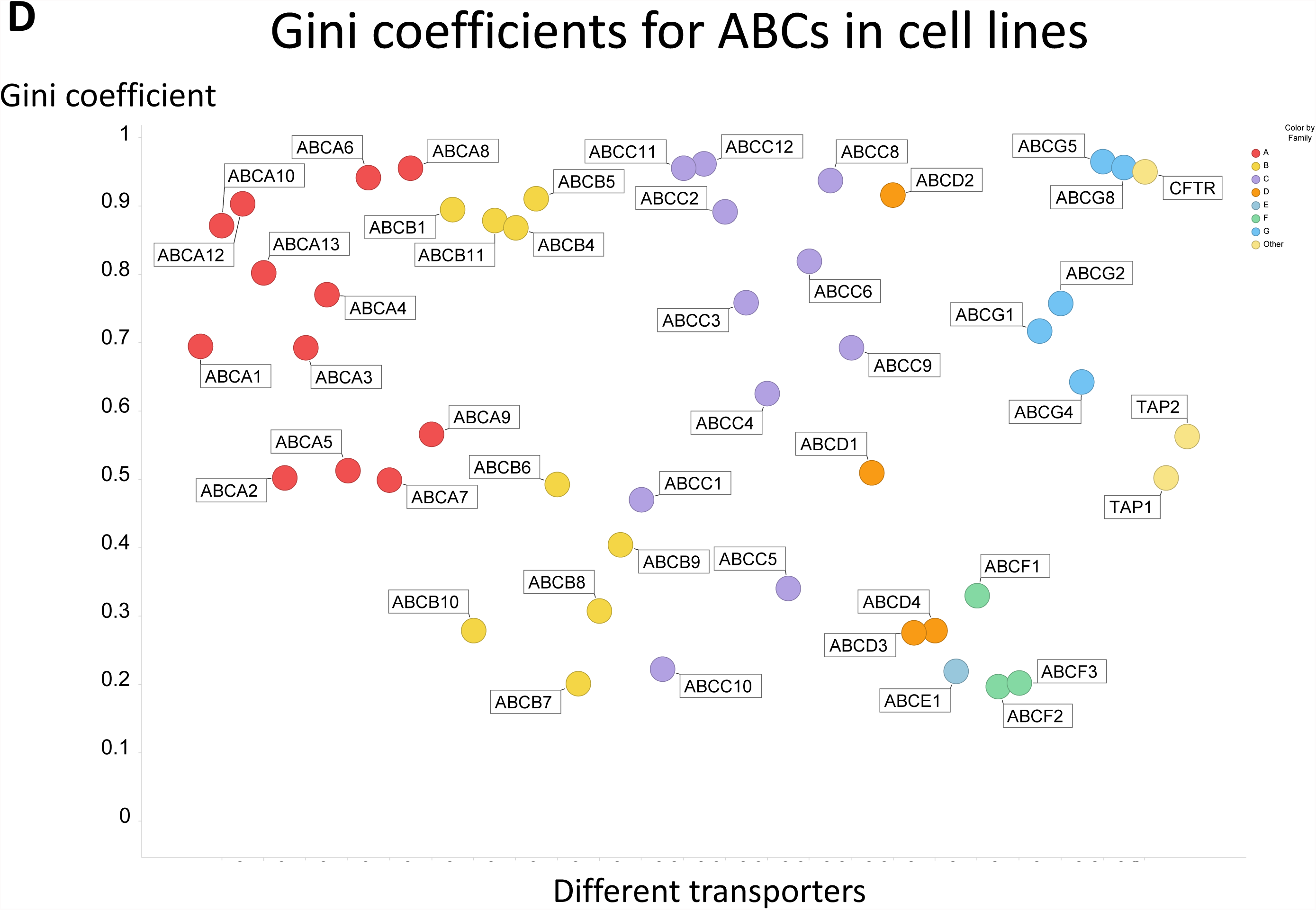
Expression profiling of various ABC transporters in 59 tissues and 56 cell lines. **A**. Minimum and maximum expression levels in the 59 tissues considered. **B**. Gini coefficient for the expression of all ABC transporters in 59 tissues. **C**. Minimum and maximum expression levels in the 56 cell lines considered. **D**. Gini coefficient for the expression of all ABC transporters in 56 cell lines.

### ABC transporters in cell lines

Figure 5C shows the minimum and maximum expression levels for all 48 ABCs, many of which lack detectable expression in at least one cell line. Again, the <u>ranges</u> of expression are considerable, and somewhat more so than those of the SLCs in tissues. No family (encoded in colour in Fig 5C) seems especially highly expressed. The overall most highly expressed ABC transporter is ABCE1. The Gini coefficients (Fig 5D) are also larger and vary more than those of both the SLCs and of the ABCs in tissues, with a median value of 0.692, suggesting adaptive selection for specialised purposes in the relevant cell lines. 11/48 GCs are greater than 0.9, while five are below 0.25. Several ABCs exhibit very high Gini coefficients that (0.964) of ABCG5 (a multi-drug resistance efflux pump) again being the largest; here it is effectively expressed only in the HepG2 cell line (a liver carcinoma cell line widely used in drug toxicity studies (Gómez-Lechón et al., 2014; Qiu et al., 2015)). By contrast, F family ABCs again tend to have lower Gini coefficients, i.e. are more universally expressed, again consistent with the view that they are not in fact transporters.

Overall, the median expression levels for SLCs are 3.27 TPM and 1.26 TPM for tissues and cell lines, respectively, while those for ABCs are 4.23 and 1.48. Thus, while many of these cell lines are cancer-derived, they do not seem to express transporters to any greater degree than to the tissues whence they originated; indeed there is a slight downregulation, consistent with earlier findings that the majority of differentially expressed genes (as transporters are) are downregulated in cancer cells (Danielsson et al., 2013).

## Overall variation in expression profiles between tissues and cells

Thus far we have focused on the differential expression of transporters per se, and paid little attention to the tissues and cell lines *per se*. This might be called the effect of tissue or cell line on expression profiles (or, as appropriate, the effect of their immortalisation thereon). The opposite dimension of a table of expression profiles focuses effectively on the opposite, viz the similarity of tissues or cell lines as judged by their expression profiles.

### Overall analysis and clustering of cell lines based on their transporter transcription levels

Although the data are far from being normally distributed, it is of interest to see which tissues and cell lines are most different from each other based solely on the expression profiles of their transporters; these data (normalised to unit variance) are given as a principal components plot in Fig 6A and B, where tissue type is encoded by colour and in the former whether it is a tumour (gray) or not is also encoded by a circular shape. Only a small amount of the variance is explained by the first two PCs, consistent with the high variability between tissues and cells, and scree plots are given as insets. (Loadings plots are correspondingly uninformative; data not shown.) However, for the cell lines a slight clustering can be observed for the lymphoid/myeloid derived cell lines as well as the telomerase immortalized cell lines. The cell line expressing the largest total amount of transporter transcripts (11,566 TPM) *in toto* is BeWo (a placental carcinoma (Orendi et al., 2010)), while that expressing the fewest (5,215 TPM) is ASC TERT1 (a human telomerase-immortalized human adipose-derived mesenchymal stem cell line (Wolbank et al., 2009)); the variance in transcripts that may be observed between these two cell lines is given in Figure 6C, with several of those with the greatest differences illustrated. Overall, the fact that the total variation in transporter expression is just twofold does show (i) the limitation of membrane ‘real estate’ area that partly controls membrane protein expression (Kell et al., 2015; Molenaar et al., 2009; Zhuang et al., 2011), and (ii) their overall importance to the cellular economy.

**Figure 6.**
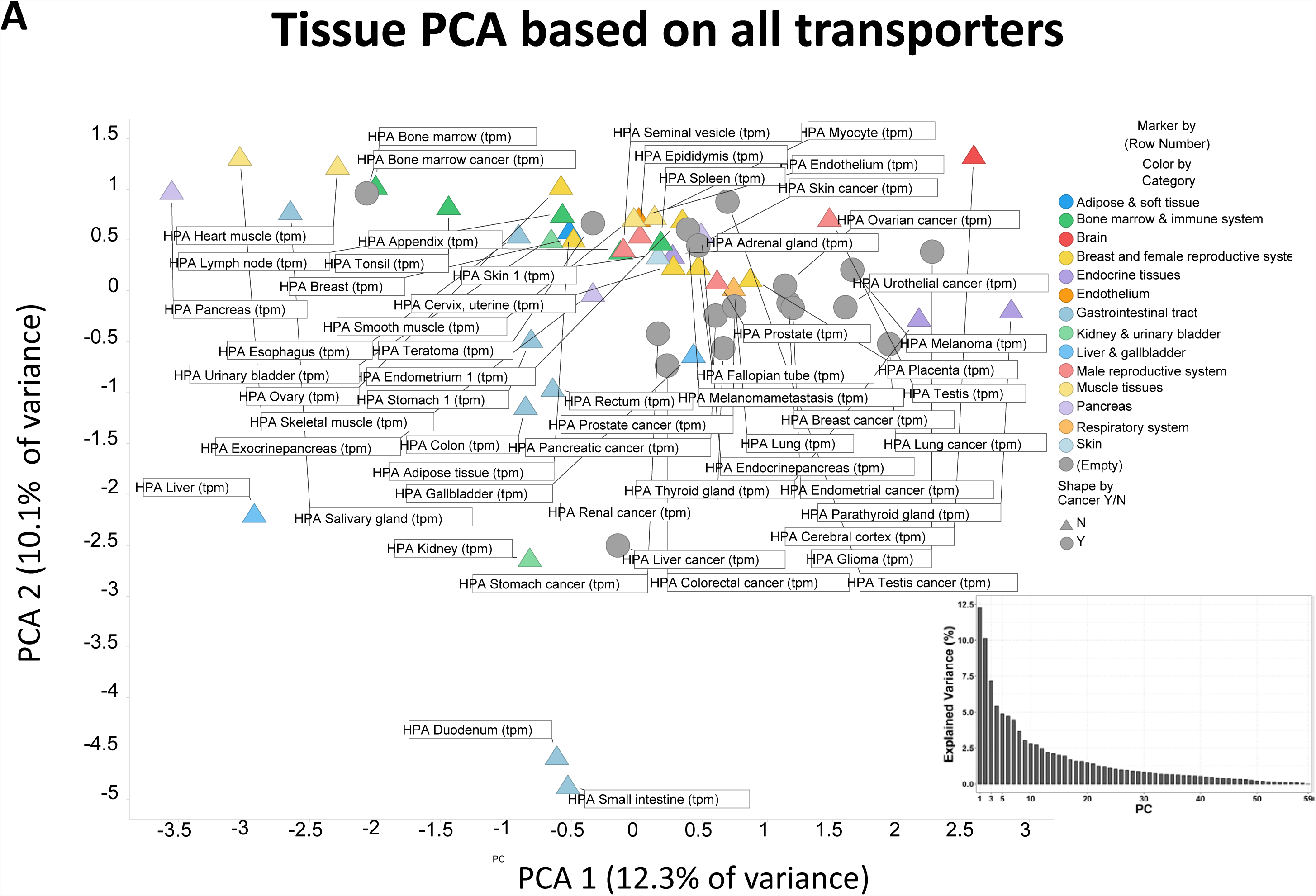
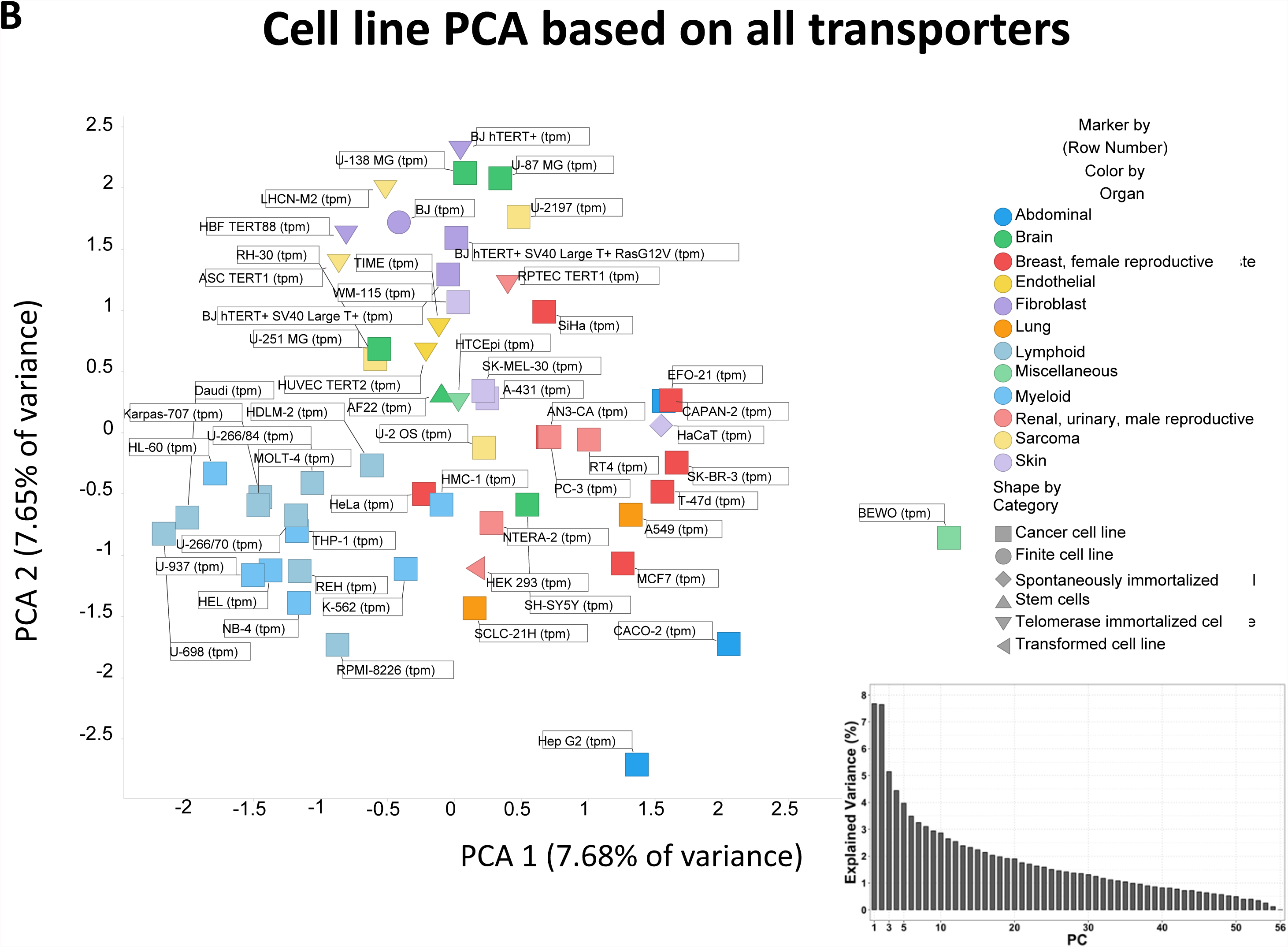
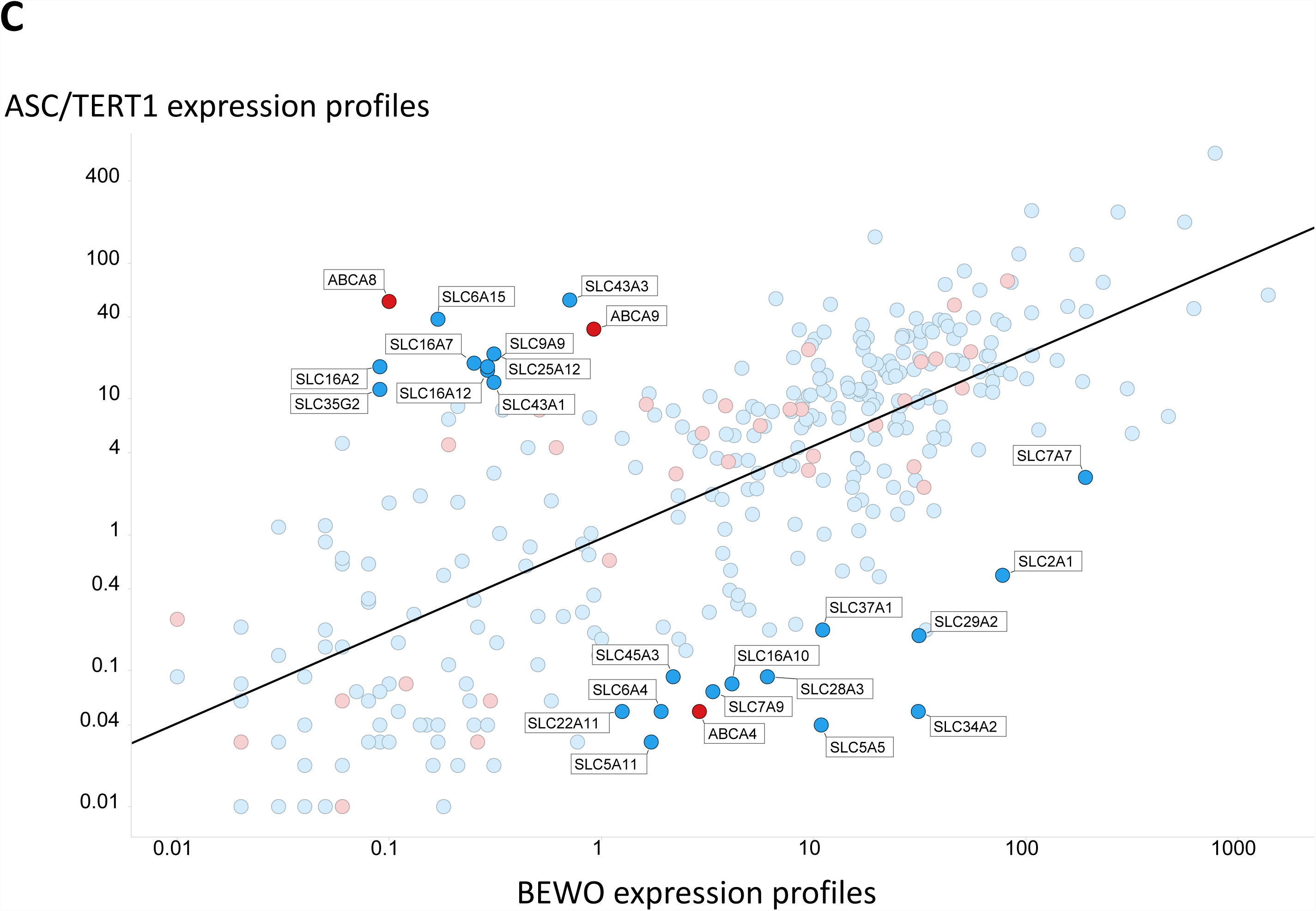
Overall variance of SLC plus ABC transporter expression in **A** different tissues and **B** different cell lines. Analyses were run in KNIME using the expression profiles of <u>both</u> SLCs and ABCs, each normalised to unit variance. Inserts in A and B represent the scree plots of %variance explained by different PCs. **C** Variance in transcript levels of both SLC (blue) and ABC (red) transporters in just two cell lines (BEWO and ASC/TERT1) (r^2^ = 0.50).

## The unusually heterogeneous nature of cell transporter expression profiles

### Tissues

We have indicated above that the values of Gini coefficient for the expression profiles of transporters between different tissues and cells tend to be unusually high, but have not yet quantified their differences relative to those of other genes (most extremely the so-called ‘housekeeping’ genes whose expression occurs in all metabolising cells and varies least). (Such genes are also typically chosen for the purposes of microbial detection and speciation (Santos and Ochman, 2004; Zeigler, 2003).) Although our focus is SLCs and ABCs versus other genes, we briefly note the differences.

From such data, the most transcribed gene over any other in cell lines is the *ATP6* gene (mitochondrial ATP synthase subunit a, Uniprot P00846, with 42,706 TPM in HeLa cells) while that in tissues is *ALB* (albumin, Uniprot P02768, with 105,947 TPM in liver). The median of all the maxima for tissues is 46 TPM, and for cell lines 40 TPM). Obviously the first of these (*ATP6* and *ALB*) are much larger numbers than those for any transporters (Figures 2, 5), but the medians (see also Fig 1B) are in quite a similar range; this again illustrates the rather specialist nature of different tissue expression profiles, including those of transporters, for different purposes.

The overall picture of the distribution of Gini coefficients between the three classes of molecule (SLC/ABC/Other) is given in Figure 7 (422 genes had very little expression at all (max = 0.25 TPM) and were ignored). Gene names are in alphabetic order, so it is clear where most of the ABCs (in blue) and SLCs (red) lie. Simply by inspection of this figure we can tell that many more ‘other’ genes (19%) have a Gini coefficient below say 0.25 than do those for SLCs (9%) and ABCs (10%). In a similar vein, 33% of SLCs and 24% of ABCs have a Gini coefficient exceeding 0.75, while 24% do for other genes. This latter high number is because of several clusters that are visible (and marked) in Fig 7A, specifically those for olfactory receptor proteins (over 300 genes, expressed in specific tissues which, given their high Gini coefficients, necessarily varied for different olfactory receptor proteins) and keratin (over 150 genes, mainly in the melanoma tissues, of which 58 are KRT for keratin and 58 KRTAP for keratin-associated proteins). Note, however, that the maximum expression level for most ORs, and for 69% of the 94 KRTAP (keratin-associated protein) genes, was mainly less than 1 TPM, and thus uncertain whether they encodes any detectable level of proteins. By contrast, transcriptional activators in the form of zinc-finger proteins (ZNF; over 500 transcripts, 82%/97% of which had a median/maximum expression greater than 1TPM) have very low Gini coefficients as they seem to play regulatory roles in almost all cells. Cyclins are of interest, as these should be expressed only in dividing cells. Thus *CCNA1*, the gene for cyclin A1 has a Gini coefficient of 0.844. However, because our focus here is on transporters, we shall not pursue all these other very interesting questions here.

**Figure 7.**
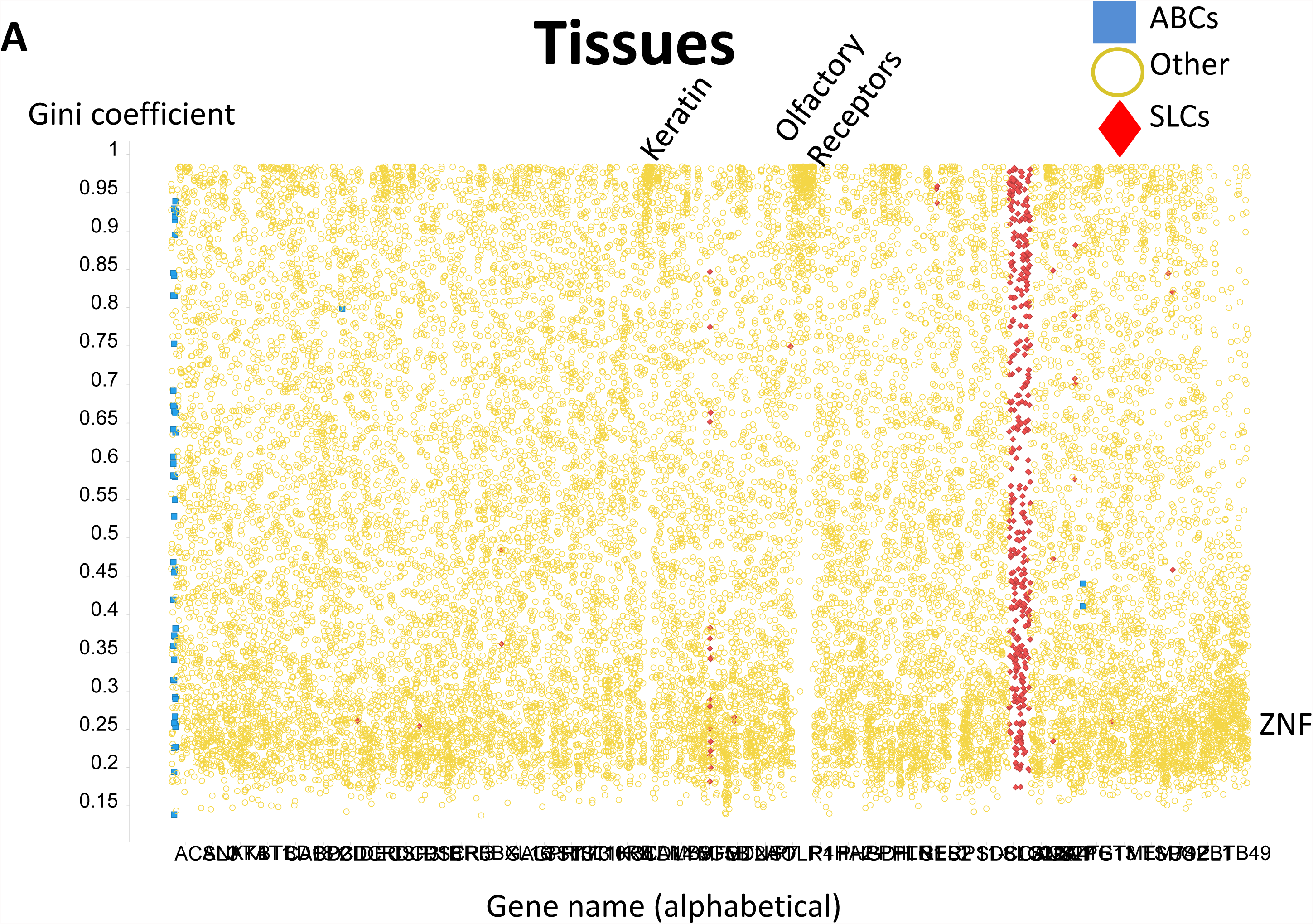
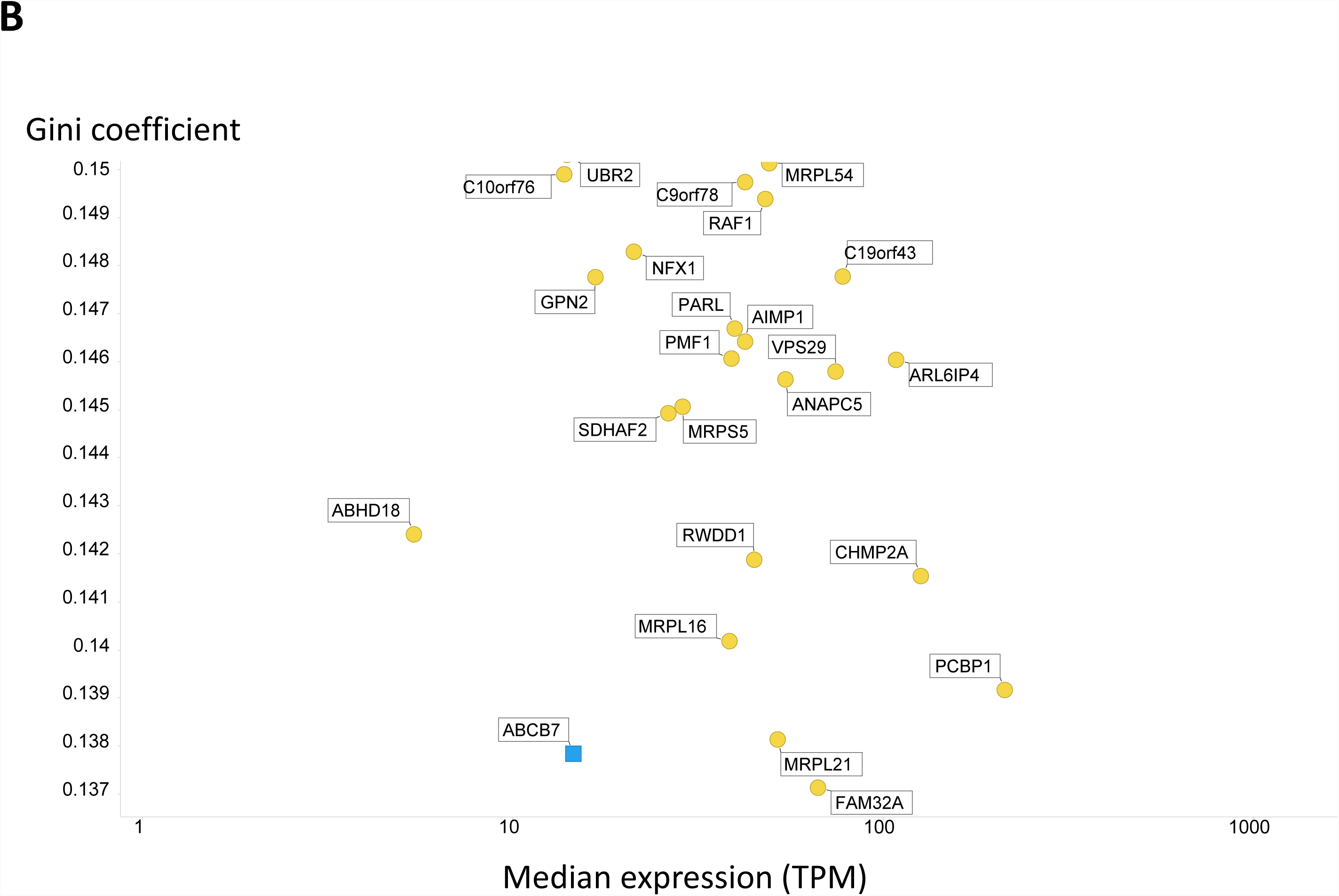
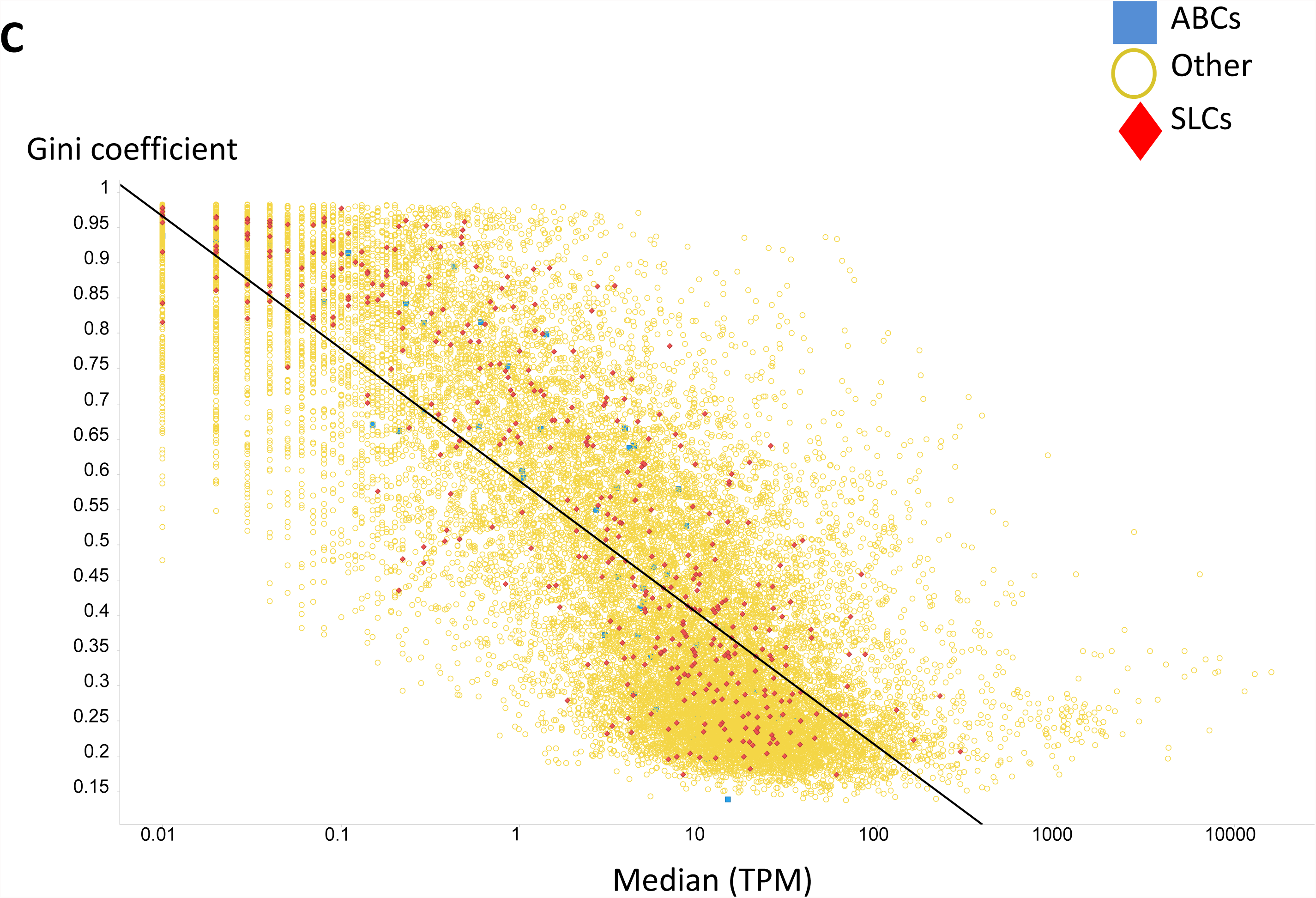
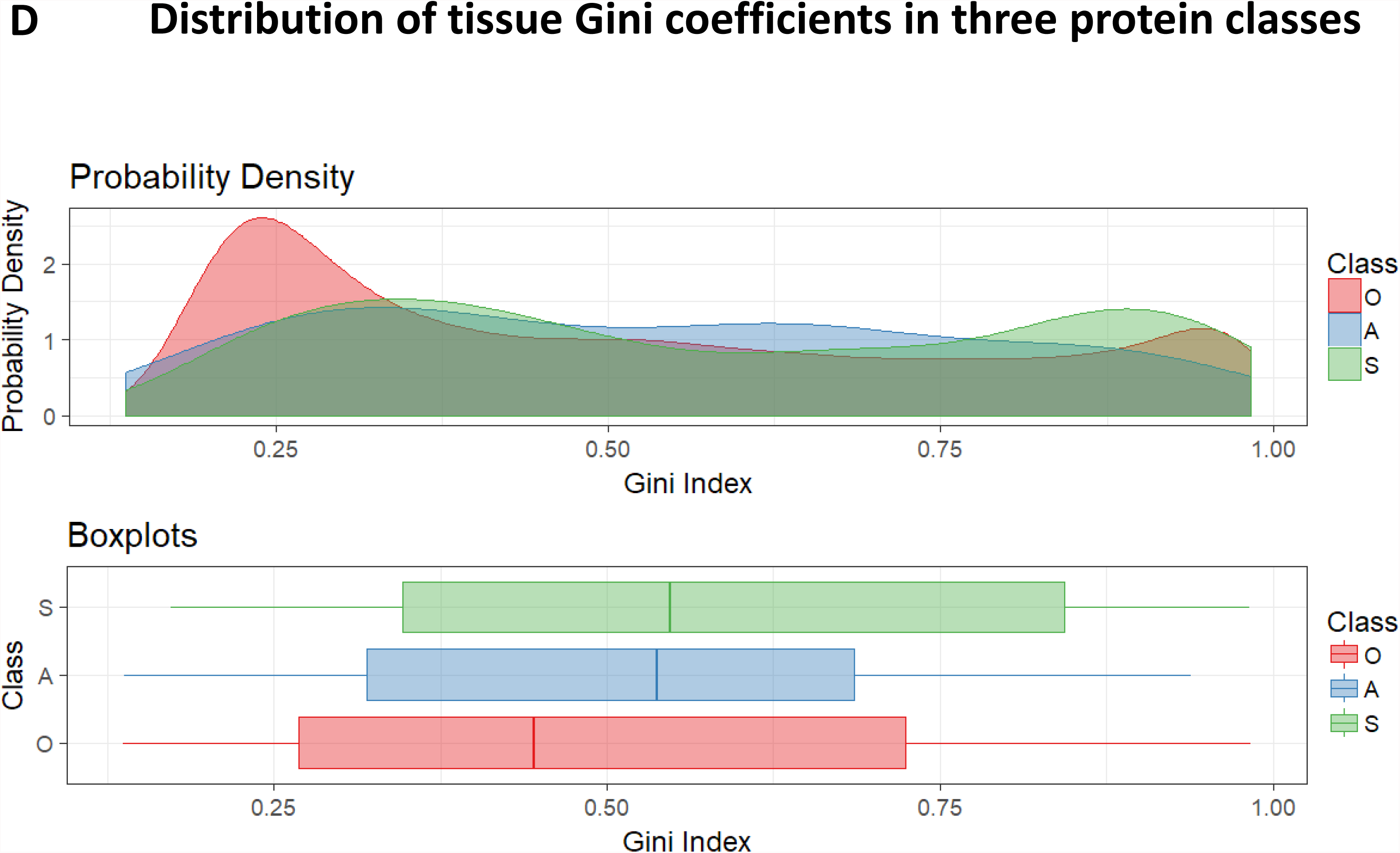
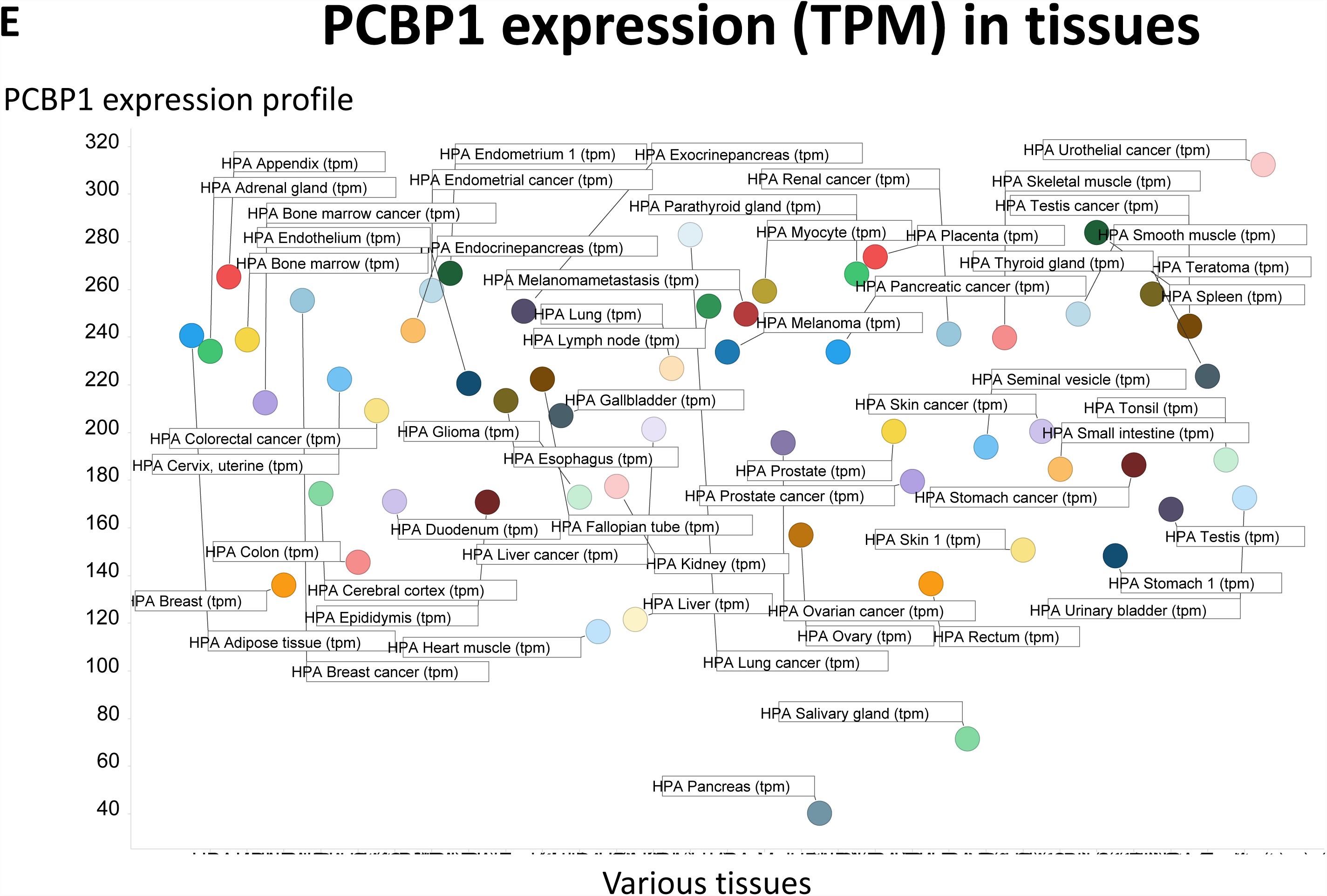
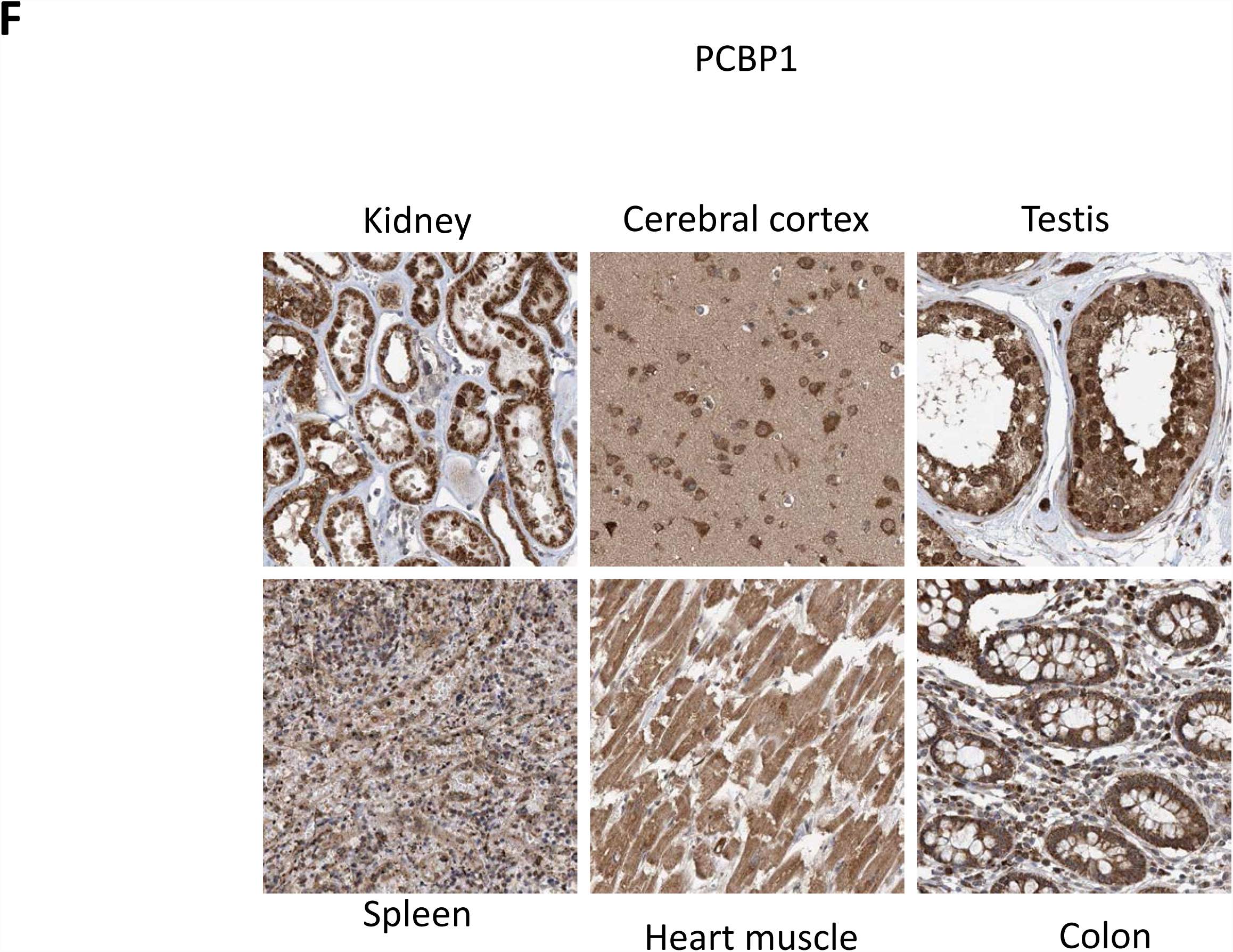
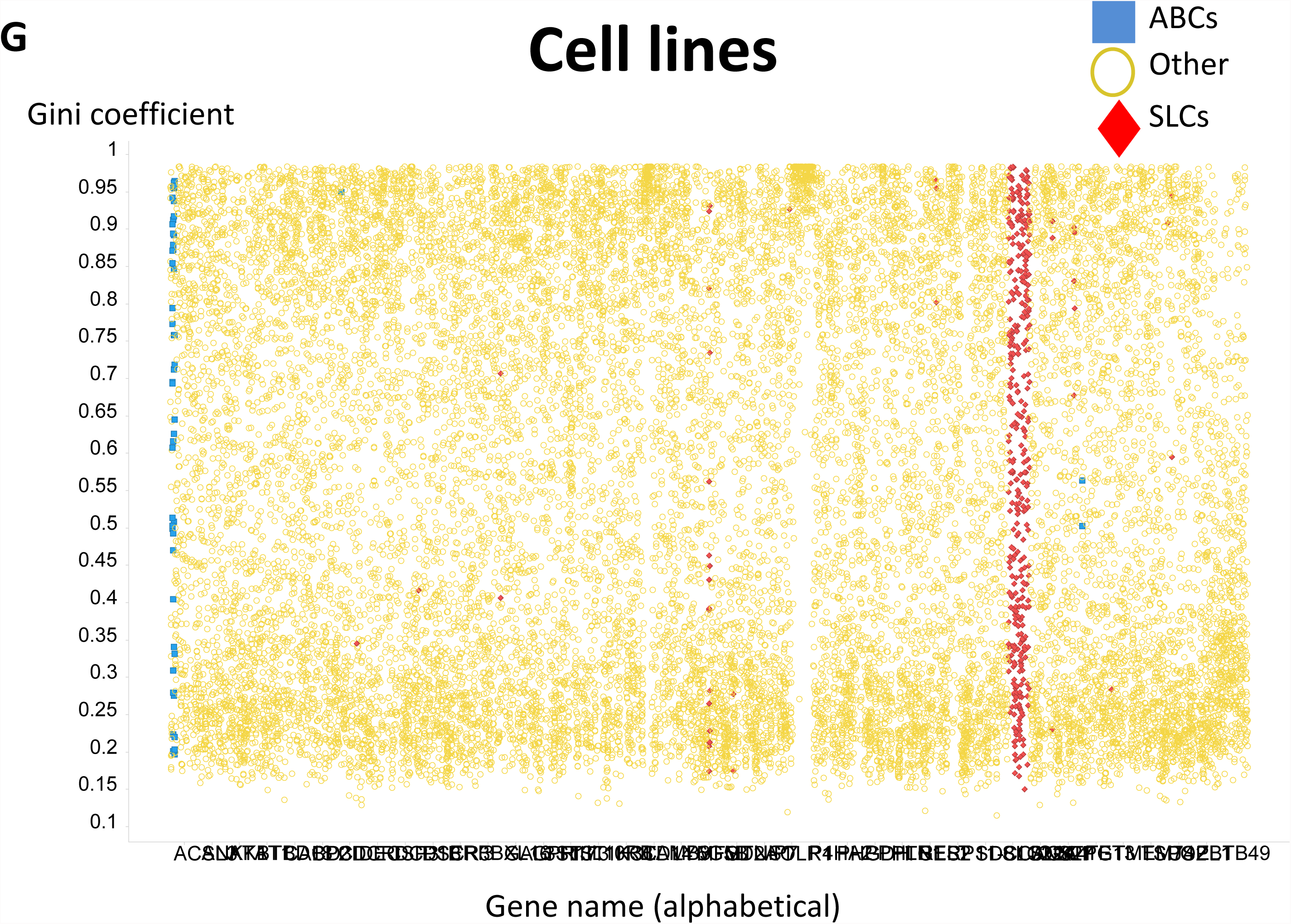
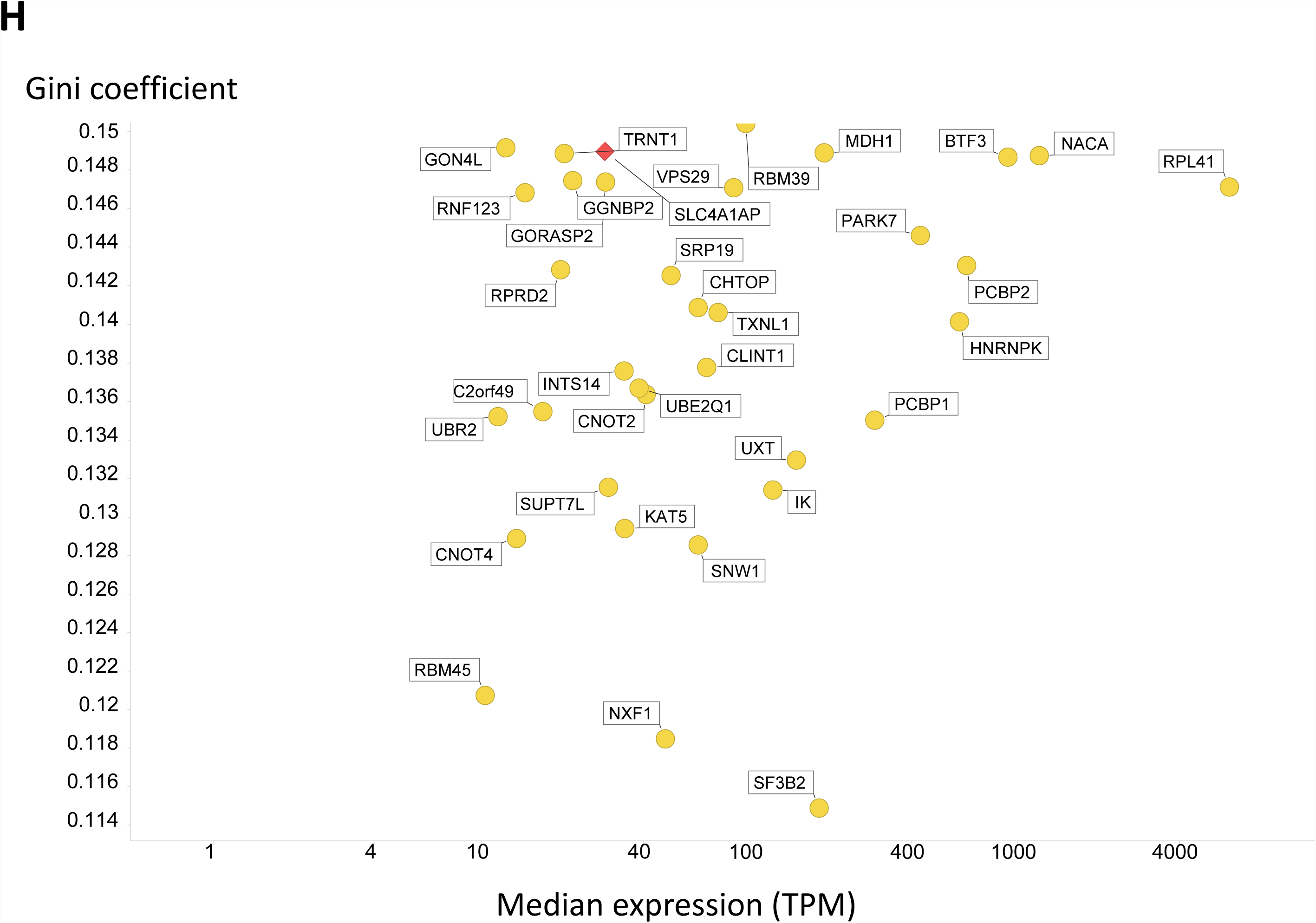
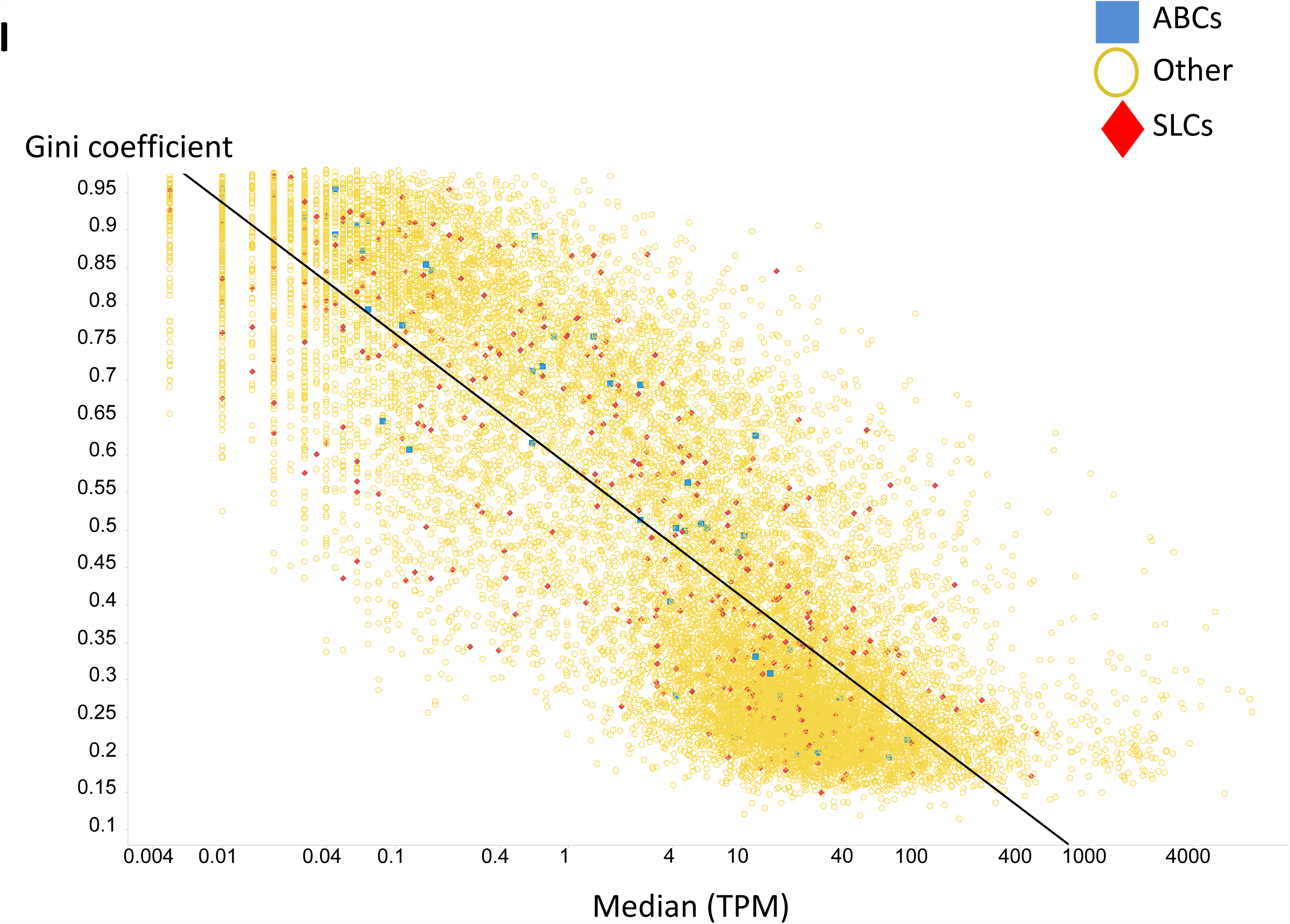
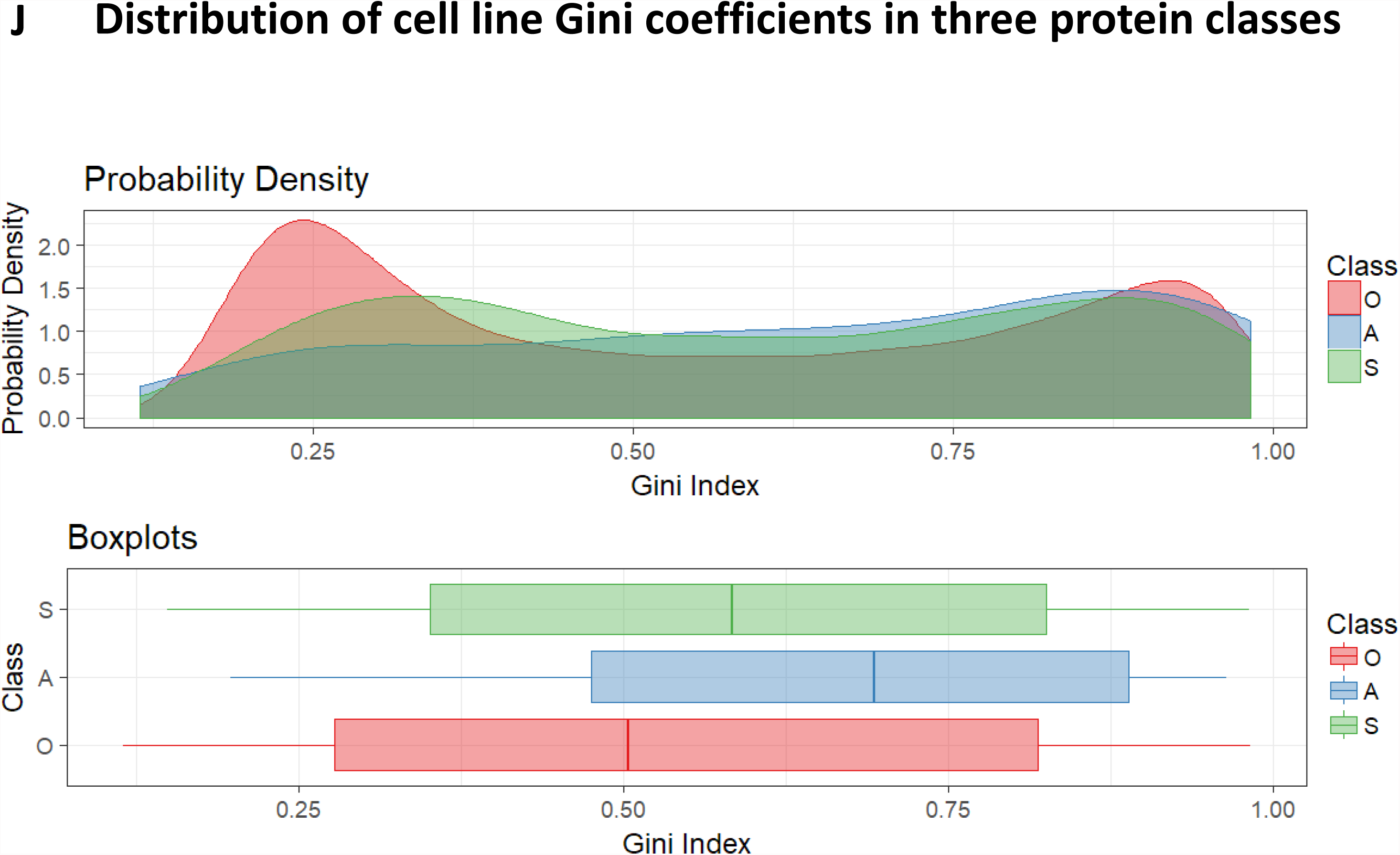
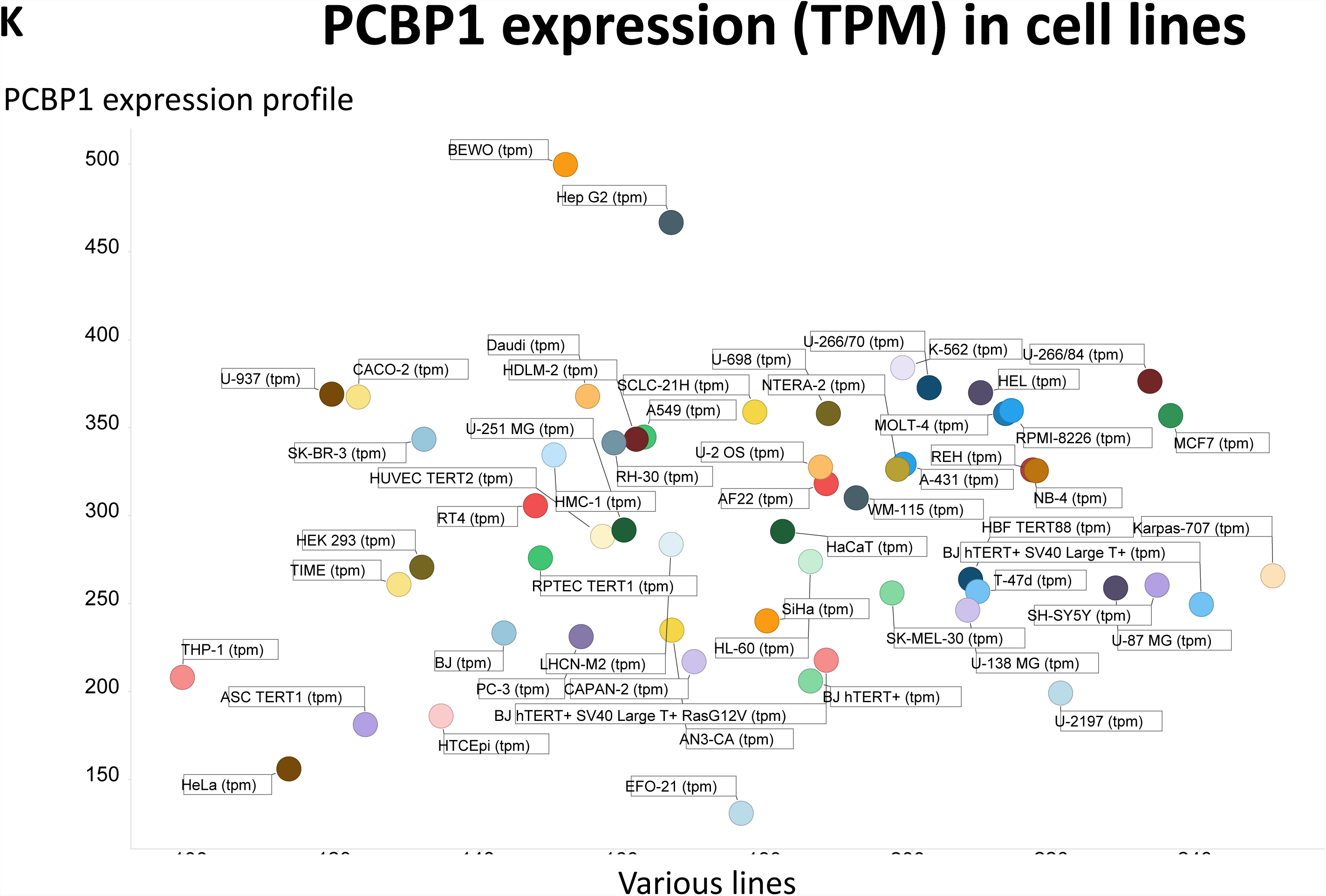
Variation of Gini coefficients of different protein classes in 59 tissues and 56 cell lines. **A**. All transcripts, alphabetically, in tissues. **B** Transcripts with a particularly low Gini coefficient in tissues. **C**. Inverse relationship between Gini index and median expression level in tissues. **D.** Distribution of Gini coefficients in the three classes of transcript in tissues. **E**. Low-Gini PCBP1 expression in tissues. **F.** Antibody-based assessment of the expression of SLC22A12 in a variety of tissues. Image edge length is 320 μm. **G.** All transcripts, alphabetically, in cell lines. **H**. Transcripts with a particularly low Gini coefficient in cell lines. **I**. Inverse relationship between Gini index and median expression level in cell lines. **J.** Distribution of Gini coefficients in the three classes in cell lines. **K**. PCBP1 expression in different cell lines.

### Genes with low expression profiles as candidate ‘housekeeping’ genes

The expression of most so-called ‘housekeeping’ genes (that are at least <u>expressed</u> in all tissues) actually varies quite widely between tissues (e.g. (de Jonge et al., 2007; Eisenberg and Levanon, 2003; Lee et al., 2002; Robinson and Oshlack, 2010)); indeed they are sufficiently different that they can be used to classify different tissues (Hsiao et al., 2001)! Here, the ‘housekeeping’ genes with the lowest Gini coefficients, hence those possibly best for normalising transcriptome or proteome experiment, are FAM32A (an RNA-binding protein; GC = 0.137), ABCB7 (a mitochondrial haem/iron exporter; GC = 0.137), MRPL16 and MRPL21 (mitoribosomal proteins; GC = 0.138), and PCBP1 (an oligo-single-stranded-dC-binding protein) GC = 0.139). Clearly their ubiquitous distribution speaks to their essentiality, and it is certainly of interest that mitoribosomal proteins have such ubiquitous expression, being somewhat equivalent to the 16S rRNA genes widely used (Caporaso et al., 2011; Cherkaoui et al., 2009; Sontakke et al., 2009; Yarza et al., 2014) in microbial taxonomy and metagenomics. Most of the other 49 large (MRPLxx) and 30 small (MRPSxx) ribosomal protein subunits also had low Gini coefficients; others with a GC of 0.15 or below are illustrated in Fig 7B, which also serves to show that most low-Gini gene products have median expression levels in the decade 20-200 TPM (so it is not a strange low-expression artefact). However, the Gini coefficients of other gene products commonly used to normalise expression profiles (e.g. (Bustin et al., 2009; de Jonge et al., 2007; Gur-Dedeoglu et al., 2009; Hoerndli et al., 2004; Li et al., 2009; Ohl et al., 2005; Oturai et al., 2016; Silver et al., 2006; Tatsumi et al., 2008; Vandesompele et al., 2002; Wang et al., 2010; Zampieri et al., 2010)) were larger, often considerably larger, e.g. *GAPDH* (glyceraldehyde 3-phosphate dehydrogenase, Uniprot P04406; 0.344), *LDHA* (lactate dehydrogenase subunit A, Uniprot P00338; 0.32), *SDHA* (succinate dehydrogenase subunit A, Uniprot P31040; 0.308), *TUBB* (tubulin β chain, Uniprot P07437; 0.333), *HPRT1* (hypoxanthine phosphoribosyl transferase 1, Uniprot P00492; 0.277), *HBS1L* (*HBS1*-like protein, Uniprot Q9Y450; 0.184), OAZ1 (ornithine decarboxylase antizyme 1, Uniprot P54368; 0.202), *PPIA* (peptidyl-prolyl cis-trans isomerase, Uniprot P62937; 0.24); *AHSP* (alpha-haemoglobin stabilising protein, Uniprot Q9NZD4; 0.97), *B2M* (β_2_-microglobulin, Uniprot P61769; 0.349), *ACTB* (β-actin, Uniprot P60709; 0.291), *HMBS* (porphobilinogen deaminase, Uniprot P08397; 0.303), *UBC* (polyubiquitin C, Uniprot P0CG48; 0.183), *POLR2F* (DNA-directed RNA polymerases I, II, and III subunit RPABC2, Uniprot P61218; 0.235), *GUSB* (β-glucuronidase, Uniprot P08236; 0.25), *TBP* (TATA-box binding protein, Uniprot P20226; 0.22) and *YWHAZ* (14-3-3 protein zeta/delta, Uniprot P63104; 0.255). However, the more recently proposed *CTBP1* (C-terminal-binding protein 1, Uniprot Q13363; 0.204) and GOLGA1 (Golgin subfamily A member 1, Uniprot Q92805; 0.189) (Lee et al., 2007) both seem like much better choices. However, the lowest Gini coefficients in tissues are FAM32A, ABCB7, MRPL21 and PCBP1 (GC 0.137 – 0.139), while the lowest three in cell lines are SF3B2, NXF1 and RBM45 (GC 0.115 – 0.122). PCBP1 is both reasonably highly expressed and has a low Gini coefficient in both tissues (0.139) and cell lines (0.135), and is an excellent novel housekeeping gene. While reference genes are often chosen to be stably expressed across variants of the same cell type rather than across different cells, this still suggests that the Gini coefficient is indeed a novel and effective way of identifying useful ‘housekeeping’ genes in expression profiling studies.

While there was no relationship between the Gini coefficient and the maximum expression (not shown), there was an interesting inverse relationship between the GC and the median expression level over all genes (Figure 7C), where the correlation coefficient was 0.62. The overall distribution of Gini coefficients for the three classes of protein (SLC/ABC/Other) is given in Figure 7D. Finally, because it was one of the gene products with the lowest Gini coefficient, as well as a reasonable expression level (median over 100 TPM), in both tissues and cell lines, we show the tissue expression profile of PCBP1 in Fig 7E; the overall variation of the great majority of these transcripts is within a two-fold range. We also illustrate its distribution in several tissues in Fig 7F.

### Cell lines

The overall data are broadly similar for cell lines (Fig 7G, although the expression of zinc fingers is less homogeneous than in the tissues). However, the genes with the lowest Gini coefficient (Fig 7H) are mostly very different from those in tissues. Note that SLC4A1AP that appears is an adaptor protein for SLC4A1 (a chloride-bicarbonate exchanger, commonly known as band 3 protein), so it is not itself a true SLC (and did not appear in Figure 4B). The very lowest, SF3B2 (Uniprot Q13435), is a subunit of an RNA splicing factor, while NXF1 (Uniprot Q9UBU9) is a nuclear export factor, and RBM45 (Uniprot Q8IUH3) RNA-binding protein 45. It is entirely reasonable that these might be expressed in all cells, and evidently at a fairly constant level. Overall, we conclude that the Gini approach is capable of finding novel housekeeping genes to act as references for microarrays and for qPCR, and will be particularly beneficial in studies employing several differentiated cell/tissue types. There is again a correlation between the Gini index and median expression level (r^2^ = 0.67) (Figure 7I). Overall, we find that 8.5% of SLCs, 16% of ABCs (including two F-family members) and 18% of other genes have a GC below 0.25, while those above 0.75 are ABC 32%, SLC 25% and Other 19%. Again, there is a significantly greater heterogeneity among transporter genes than among other genes when taken as a whole (Fig 7I). Finally, Fig 7J shows the expression profile of PCBP1 in cell lines; again the overwhelming majority are within a two-fold range.

## Discussion and conclusions

The present paper has highlighted at least three main areas. First, we exploit the Gini coefficient as a novel, convenient and easily understandable metric for reflecting how unequally a given transcript is expressed in a series of tissues or cell lines. In contrast to its usual use in economics, where it ranges from ~0.25 to ~0.51 in different countries, the Gini index here ranged from as low as 0.11 to as high as 0.98, reflecting in the latter case virtually unique expression in particular tissues. In many cases, the biology underpinning this is quite opaque, but the purpose of data-driven studies is to generate rather than to test hypotheses (Kell and Oliver, 2004). We also recognise here that we paid relatively little attention to the distribution of transporters <u>within</u> different tissues and their potential cell-type-specific distribution within an organ (e.g. (Bahar Halpern et al., 2017)), where they presumably account for the very striking intra-organ distributions of drugs (e.g. (Römpp et al., 2011)); that will have to be a subject for further work.

A second chief area of interest is the distribution of transporters between different tissues. A detailed analysis showed that they tended to have significantly higher Gini coefficients than did other genes. This illustrates the point that despite the fact that their substrates are almost uniformly available via the bloodstream, and biochemistry textbooks and wallcharts largely show this, they clearly use substrates differentially (ergothioneine and the SLC22A4 transporter being a nice example (Gründemann, 2012; Gründemann et al., 2005)). It also implies strongly that in many cases we do not in fact know the natural substrates, many of which are clearly exogenous (O’Hagan and Kell, 2017b).

The third main recognition is that the Gini index provides a particularly useful, convenient, non-parametric and intelligible means of identifying those genes whose expression profile varies least across a series of cells or tissues, thus providing a novel and convenient strategy for the identification of those housekeeping genes best used as genes against which to normalise other expression profiles in a variety of studies. We have here highlighted quite a number that have not previously been so identified.

Overall, we consider that assessing the Gini index for the distribution of particular transporters and other proteins between different cells has much to offer the development of novel biology; it should prove a highly useful addition to the armoury of both the systems biologist and the data analyst.

## Acknowledgments

DBK and PJD thank the BBSRC (grant BB/P009042/1) for financial support. EL thanks the entire staff of the Human Protein Atlas program. EL acknowledges financial support by the Knut and Alice Wallenberg Foundation, the Erling Persson Foundation and facility support to the Science for Life Laboratory (SciLifeLab).

## References

Ainali, C., Valeyev, N., Perera, G., Williams, A., Gudjonsson, J.E., Ouzounis, C.A., Nestle, F.O., and Tsoka, S. (2012). Transcriptome classification reveals molecular subtypes in psoriasis. BMC Genomics 13, 472.

Almén, M.S., Nordström, K.J., Fredriksson, R., and Schiöth, H.B. (2009). Mapping the human membrane proteome: a majority of the human membrane proteins can be classified according to function and evolutionary origin. BMC biology 7, 50.

Anzai, N., Jutabha, P., Amonpatumrat-Takahashi, S., and Sakurai, H. (2012). Recent advances in renal urate transport: characterization of candidate transporters indicated by genome-wide association studies. Clin Exp Nephrol 16, 89–95.

Bahar Halpern, K., Shenhav, R., Matcovitch-Natan, O., Toth, B., Lemze, D., Golan, M., Massasa, E.E., Baydatch, S., Landen, S., Moor, A.E., et al. (2017). Single-cell spatial reconstruction reveals global division of labour in the mammalian liver. Nature 542, 352–356.

Baker, S.G. (2003). The central role of receiver operating characteristic (ROC) curves in evaluating tests for the early detection of cancer. J Natl Cancer Inst 95, 511–515.

Berthold, M.R., Cebron, N., Dill, F., Gabriel, T.R., Kötter, T., Meinl, T., Ohl, P., Sieb, C., Thiel, K., and Wiswedel, B. (2008). KNIME: the Konstanz Information Miner. In Data Analysis, Machine Learning and Applications, C. Preisach, H. Burkhardt, L. Schmidt-Thieme, and R. Decker, eds. (Berlin, Springer), pp. 319–326.

Bray, N.L., Pimentel, H., Melsted, P., and Pachter, L. (2016). Near-optimal probabilistic RNA-seq quantification. Nat Biotechnol 34, 525–527.

Broadhurst, D., and Kell, D.B. (2006). Statistical strategies for avoiding false discoveries in metabolomics and related experiments. Metabolomics 2, 171–196.

Bröer, S., and Palacín, M. (2011). The role of amino acid transporters in inherited and acquired diseases. Biochem J 436, 193–211.

Brower, J.V., Lim, C.H., Jorgensen, M., Oh, S.P., and Terada, N. (2009). Adenine nucleotide translocase 4 deficiency leads to early meiotic arrest of murine male germ cells. Reproduction 138, 463–470.

Brower, J.V., Rodic, N., Seki, T., Jorgensen, M., Fliess, N., Yachnis, A.T., McCarrey, J.R., Oh, S.P., and Terada, N. (2007). Evolutionarily conserved mammalian adenine nucleotide translocase 4 is essential for spermatogenesis. J Biol Chem 282, 29658–29666.

Bustin, S.A., Benes, V., Garson, J.A., Hellemans, J., Huggett, J., Kubista, M., Mueller, R., Nolan, T., Pfaffl, M.W., Shipley, G.L., et al. (2009). The MIQE guidelines: minimum information for publication of quantitative real-time PCR experiments. Clin Chem 55, 611–622.

Caporaso, J.G., Lauber, C.L., Walters, W.A., Berg-Lyons, D., Lozupone, C.A., Turnbaugh, P.J., Fierer, N., and Knight, R. (2011). Global patterns of 16S rRNA diversity at a depth of millions of sequences per sample. Proc Natl Acad Sci U S A 108 Suppl 1, 4516–4522.

Ceriani, L., and Verme, P. (2012). The origins of the Gini index: extracts from Variabilità e Mutabilità (1912) by Corrado Gini. J Econ Inequal 10, 421–443.

César-Razquin, A., Snijder, B., Frappier-Brinton, T., Isserlin, R., Gyimesi, G., Bai, X., Reithmeier, R.A., Hepworth, D., Hediger, M.A., Edwards, A.M., et al. (2015). A call for systematic research on solute carriers. Cell 162, 478–487.

Chen, Z., Shi, T., Zhang, L., Zhu, P., Deng, M., Huang, C., Hu, T., Jiang, L., and Li, J. (2016). Mammalian drug efflux transporters of the ATP binding cassette (ABC) family in multidrug resistance: A review of the past decade. Cancer Lett 370, 153–164.

Cherkaoui, A., Emonet, S., Ceroni, D., Candolfi, B., Hibbs, J., Francois, P., and Schrenzel, J. (2009). Development and validation of a modified broad-range 16S rDNA PCR for diagnostic purposes in clinical microbiology. J Microbiol Methods 79, 227–231.

Clémençon, B., Babot, M., and Trezeguet, V. (2013). The mitochondrial ADP/ATP carrier (SLC25 family): Pathological implications of its dysfunction. Mol Aspects Med 34, 485–493.

Colas, C., Ung, P.M.U., and Schlessinger, A. (2016). SLC transporters: structure, function, and drug discovery. MedChemComm 7, 1069–1081.

da Cunha, J.P.C., Galante, P.A., de Souza, J.E., de Souza, R.F., Carvalho, P.M., Ohara, D.T., Moura, R.P., Oba-Shinja, S.M., Marie, S.K., Silva, W.A., Jr., et al. (2009). Bioinformatics construction of the human cell surfaceome. Proc Natl Acad Sci U S A 106, 16752–16757.

Damgaard, C., and Weiner, J. (2000). Describing inequality in plant size or fecundity. Ecology 81, 1139–1142.

Danielsson, F., Skogs, M., Huss, M., Rexhepaj, E., O’Hurley, G., Klevebring, D., Pontén, F., Gad, A.K.B., Uhlén, M., and Lundberg, E. (2013). Majority of differentially expressed genes are down-regulated during malignant transformation in a four-stage model. Proc Natl Acad Sci U S A 110, 6853–6858.

de Jonge, H.J.M., Fehrmann, R.S.N., de Bont, E.S.J.M., Hofstra, R.M.W., Gerbens, F., Kamps, W.A., de Vries, E.G.E., van der Zee, A.G.J., te Meerman, G.J., and ter Elst, A. (2007). Evidence based selection of housekeeping genes. PLoS One 2, e898.

Dean, M., and Annilo, T. (2005). Evolution of the ATP-binding cassette (ABC) transporter superfamily in vertebrates. Annu Rev Genomics Hum Genet 6, 123–142.

Dobson, P.D., and Kell, D.B. (2008). Carrier-mediated cellular uptake of pharmaceutical drugs: an exception or the rule? Nat Rev Drug Disc 7, 205–220.

Dolce, V., Scarcia, P., Iacopetta, D., and Palmieri, F. (2005). A fourth ADP/ATP carrier isoform in man: identification, bacterial expression, functional characterization and tissue distribution. FEBS Lett 579, 633–637.

Eadie, L.N., Hughes, T.P., and White, D.L. (2014). Interaction of the efflux transporters ABCB1 and ABCG2 with imatinib, nilotinib, and dasatinib. Clin Pharmacol Ther 95, 294–306.

Eisen, M.B., Spellman, P.T., Brown, P.O., and Botstein, D. (1998). Cluster analysis and display of genome-wide expression patterns. Proc Natl Acad Sci 95, 14863–14868.

Eisenberg, E., and Levanon, E.Y. (2003). Human housekeeping genes are compact. Trends Genet 19, 362–365.

Fagerberg, L., Hallström, B.M., Oksvold, P., Kampf, C., Djureinovic, D., Odeberg, J., Habuka, M., Tahmasebpoor, S., Danielsson, A., Edlund, K., et al. (2014). Analysis of the human tissue-specific expression by genome-wide integration of transcriptomics and antibody-based proteomics. Mol Cell Proteomics 13, 397–406.

Fredriksson, R., Nordström, K.J., Stephansson, O., Hägglund, M.G., and Schiöth, H.B. (2008). The solute carrier (SLC) complement of the human genome: phylogenetic classification reveals four major families. FEBS Lett 582, 3811–3816.

Giacomini, K.M., and Huang, S.M. (2013). Transporters in drug development and clinical pharmacology. Clin Pharmacol Ther 94, 3–9.

Giacomini, K.M., Huang, S.M., Tweedie, D.J., Benet, L.Z., Brouwer, K.L., Chu, X., Dahlin, A., Evers, R., Fischer, V., Hillgren, K.M., et al. (2010). Membrane transporters in drug development. Nat Rev Drug Discov 9, 215–236.

Gini, C. (1909). Concentration and dependency ratios (in Italian). English translation in: Rivista di Politica Economica, 87 (1997), 769–789.

Gini, C. (1912). Variabilità e Mutabilità. Contributo allo Studio delle Distribuzioni e delle Relazioni Statistiche (Bologna, C. Cuppini).

Gómez-Lechón, M.J., Tolosa, L., and Donato, M.T. (2014). Cell-based models to predict human hepatotoxicity of drugs. Rev Toxicol 31, 149–156.

Grixti, J., O’Hagan, S., Day, P.J., and Kell, D.B. (2017). Enhancing drug efficacy and therapeutic index through cheminformatics-based selection of small molecule binary weapons that improve transporter-mediated targeting: a cytotoxicity system based on gemcitabine. Front Pharmacol 8, 155.

Gründemann, D. (2012). The ergothioneine transporter controls and indicates ergothioneine activity--a review. Prev Med 54 Suppl, S71–74.

Gründemann, D., Harlfinger, S., Golz, S., Geerts, A., Lazar, A., Berkels, R., Jung, N., Rubbert, A., and Schömig, E. (2005). Discovery of the ergothioneine transporter. Proc Natl Acad Sci U S A 102, 5256–5261.

Gur-Dedeoglu, B., Konu, O., Bozkurt, B., Ergul, G., Seckin, S., and Yulug, I.G. (2009). Identification of endogenous reference genes for qRT-PCR analysis in normal matched breast tumor tissues. Oncol Res 17, 353–365.

Hagenbuch, B., and Stieger, B. (2013). The SLCO (former SLC21) superfamily of transporters. Mol Aspects Med 34, 396–412.

Hamazaki, T., Leung, W.Y., Cain, B.D., Ostrov, D.A., Thorsness, P.E., and Terada, N. (2011). Functional expression of human adenine nucleotide translocase 4 in Saccharomyces cerevisiae. PLoS One 6, e19250.

Hediger, M.A., Clemencon, B., Burrier, R.E., and Bruford, E.A. (2013). The ABCs of membrane transporters in health and disease (SLC series): Introduction. Mol Aspects Med 34, 95–107.

Hoerndli, F.J., Toigo, M., Schild, A., Götz, J., and Day, P.J. (2004). Reference genes identified in SH-SY5Y cells using custom-made gene arrays with validation by quantitative polymerase chain reaction. Anal Biochem 335, 30–41.

Höglund, P.J., Nordström, K.J.V., Schiöth, H.B., and Fredriksson, R. (2011). The solute carrier families have a remarkably long evolutionary history with the majority of the human families present before divergence of Bilaterian species. Mol Biol Evol 28, 1531–1541.

Hsiao, L.L., Dangond, F., Yoshida, T., Hong, R., Jensen, R.V., Misra, J., Dillon, W., Lee, K.F., Clark, K.E., Haverty, P., et al. (2001). A compendium of gene expression in normal human tissues. Physiol Genomics 7, 97–104.

Jack, D.L., Paulsen, I.T., and Saier, M.H. (2000). The amino acid/polyamine/organocation (APC) superfamily of transporters specific for amino acids, polyamines and organocations. Microbiology 146 (Pt 8), 1797–1814.

Jeong, J., and Eide, D.J. (2013). The SLC39 family of zinc transporters. Mol Aspects Med 34, 612–619.

Jiang, L., Chen, H., Pinello, L., and Yuan, G.C. (2016). GiniClust: detecting rare cell types from single-cell gene expression data with Gini index. Genome Biol 17, 144.

Kell, D.B. (2015). The transporter-mediated cellular uptake of pharmaceutical drugs is based on their metabolite-likeness and not on their bulk biophysical properties: Towards a systems pharmacology Perspect Sci 6, 66–83.

Kell, D.B. (2016). How drugs pass through biological cell membranes – a paradigm shift in our understanding? Beilstein Magazine 2, http://www.beilstein-institut.de/download/628/609_kell.pdf.

Kell, D.B., Dobson, P.D., Bilsland, E., and Oliver, S.G. (2013). The promiscuous binding of pharmaceutical drugs and their transporter-mediated uptake into cells: what we (need to) know and how we can do so. Drug Disc Today 18, 218–239.

Kell, D.B., Dobson, P.D., and Oliver, S.G. (2011). Pharmaceutical drug transport: the issues and the implications that it is essentially carrier-mediated only. Drug Disc Today 16, 704–714.

Kell, D.B., and Oliver, S.G. (2004). Here is the evidence, now what is the hypothesis? The complementary roles of inductive and hypothesis-driven science in the post-genomic era. Bioessays 26, 99–105.

Kell, D.B., and Oliver, S.G. (2014). How drugs get into cells: tested and testable predictions to help discriminate between transporter-mediated uptake and lipoidal bilayer diffusion. Front Pharmacol 5, 231.

Kell, D.B., Swainston, N., Pir, P., and Oliver, S.G. (2015). Membrane transporter engineering in industrial biotechnology and whole-cell biocatalysis. Trends Biotechnol 33, 237–246.

Koepsell, H. (2013). The SLC22 family with transporters of organic cations, anions and zwitterions. Mol Aspects Med 34, 413–435.

Kohli, M.A., Lucae, S., Saemann, P.G., Schmidt, M.V., Demirkan, A., Hek, K., Czamara, D., Alexander, M., Salyakina, D., Ripke, S., et al. (2011). The neuronal transporter gene SLC6A15 confers risk to major depression. Neuron 70, 252–265.

Kondo, N., van Dam, R.M., Sembajwe, G., Subramanian, S.V., Kawachi, I., and Yamagata, Z. (2012). Income inequality and health: the role of population size, inequality threshold, period effects and lag effects. J Epidemiol Community Health 66, e11.

Lee, P.D., Sladek, R., Greenwood, C.M., and Hudson, T.J. (2002). Control genes and variability: absence of ubiquitous reference transcripts in diverse mammalian expression studies. Genome Res 12, 292–297.

Lee, S., Jo, M., Lee, J., Koh, S.S., and Kim, S. (2007). Identification of novel universal housekeeping genes by statistical analysis of microarray data. J Biochem Mol Biol 40, 226–231.

Lee, W.C. (1999). Probabilistic analysis of global performances of diagnostic tests: interpreting the Lorenz curve-based summary measures. Stat Med 18, 455–471.

Li, Y.L., Ye, F., Hu, Y., Lu, W.G., and Xie, X. (2009). Identification of suitable reference genes for gene expression studies of human serous ovarian cancer by real-time polymerase chain reaction. Anal Biochem 394, 110–116.

Lin, L., Yee, S.W., Kim, R.B., and Giacomini, K.M. (2015). SLC transporters as therapeutic targets: emerging opportunities. Nat Rev Drug Discov 14, 543–560.

Linden, A. (2006). Measuring diagnostic and predictive accuracy in disease management: an introduction to receiver operating characteristic (ROC) analysis. J Eval Clin Pract 12, 132–139.

Masters, J.R.W. (2000). Human cancer cell lines: fact and fantasy. Nat Rev Mol Cell Biol 1, 233–236.

Molenaar, D., van Berlo, R., de Ridder, D., and Teusink, B. (2009). Shifts in growth strategies reflect tradeoffs in cellular economics. Mol Syst Biol 5, 323.

Montanari, F., and Ecker, G.F. (2015). Prediction of Drug-ABC Transporter Interaction - Recent Advances and Future Challenges. Adv Drug Deliv Rev.

Mueckler, M., and Thorens, B. (2013). The SLC2 (GLUT) family of membrane transporters. Mol Aspects Med 34, 121–138.

Nishimura, S., Tsuda, H., Ito, K., Jobo, T., Yaegashi, N., Inoue, T., Sudo, T., Berkowitz, R.S., and Mok, S.C. (2007). Differential expression of ABCF2 protein among different histologic types of epithelial ovarian cancer and in clear cell adenocarcinomas of different organs. Hum Pathol 38, 134–139.

O’Hagan, S., and Kell, D.B. (2017a). Analysis of drug-endogenous human metabolite similarities in terms of their maximum common substructures. J Cheminform 9, 18.

O’Hagan, S., and Kell, D.B. (2017b). Consensus rank orderings of molecular fingerprints illustrate the ‘most genuine’ similarities between marketed drugs and small endogenous human metabolites, but highlight exogenous natural products as the most important ‘natural’ drug transporter substrates. ADMET & DMPK 5, 85–125.

O’Hagan, S., and Kell, D.B. (2017c). Consensus rank orderings of molecular fingerprints illustrate the ‘most genuine’ similarities between marketed drugs and small endogenous human metabolites, but highlight exogenous natural products as the most important ‘natural’ drug transporter substrates. bioRxiv version. bioRxiv, 110437.

O’Hagan, S., Swainston, N., Handl, J., and Kell, D.B. (2015). A ‘rule of 0.5′ for the metabolite-likeness of approved pharmaceutical drugs. Metabolomics 11, 323–339.

O’Hagan, S., and Kell, D.B. (2017). Consensus rank orderings of molecular fingerprints illustrate the ‘most genuine’ similarities between marketed drugs and small endogenous human metabolites, but highlight exogenous natural products as the most important ‘natural’ drug transporter substrates. ADME & DMPK, in the press.

Ohl, F., Jung, M., Xu, C., Stephan, C., Rabien, A., Burkhardt, M., Nitsche, A., Kristiansen, G., Loening, S.A., Radonic, A., et al. (2005). Gene expression studies in prostate cancer tissue: which reference gene should be selected for normalization? J Mol Med (Berl) 83, 1014–1024.

Orendi, K., Gauster, M., Moser, G., Meiri, H., and Huppertz, B. (2010). The choriocarcinoma cell line BeWo: syncytial fusion and expression of syncytium-specific proteins. Reproduction 140, 759–766.

Oturai, D.B., Sondergaard, H.B., Bornsen, L., Sellebjerg, F., and Christensen, J.R. (2016). Identification of Suitable Reference Genes for Peripheral Blood Mononuclear Cell Subset Studies in Multiple Sclerosis. Scand J Immunol 83, 72–80.

Palm, W., and Thompson, C.B. (2017). Nutrient acquisition strategies of mammalian cells. Nature 546, 234–242.

Palmieri, F. (2013). The mitochondrial transporter family SLC25: Identification, properties and physiopathology. Molecular aspects of medicine 34, 465–484.

Pelis, R.M., and Wright, S.H. (2014). SLC22, SLC44, and SLC47 transporters--organic anion and cation transporters: molecular and cellular properties. Curr Top Membr 73, 233–261.

Perland, E., and Fredriksson, R. (2017). Classification Systems of Secondary Active Transporters. Trends Pharmacol Sci 38, 305–315.

Perland, E., Lekholm, E., Eriksson, M.M., Bagchi, S., Arapi, V., and Fredriksson, R. (2016). The Putative SLC Transporters Mfsd5 and Mfsd11 Are Abundantly Expressed in the Mouse Brain and Have a Potential Role in Energy Homeostasis. PLoS One 11, e0156912.

Pfefferkorn, J.A. (2013). Strategies for the design of hepatoselective glucokinase activators to treat type 2 diabetes. Expert Opin Drug Discov 8, 319–330.

Pfefferkorn, J.A., Guzman-Perez, A., Litchfield, J., Aiello, R., Treadway, J.L., Pettersen, J., Minich, M.L., Filipski, K.J., Jones, C.S., Tu, M., et al. (2012). Discovery of (S)-6-(3-cyclopentyl-2-(4-(trifluoromethyl)-1H-imidazol-1-yl)propanamido)nicotini c acid as a hepatoselective glucokinase activator clinical candidate for treating type 2 diabetes mellitus. J Med Chem 55, 1318–1333.

Pfefferkorn, J.A., Litchfield, J., Hutchings, R., Cheng, X.M., Larsen, S.D., Auerbach, B., Bush, M.R., Lee, C., Erasga, N., Bowles, D.M., et al. (2011). Discovery of novel hepatoselective HMG-CoA reductase inhibitors for treating hypercholesterolemia: A bench-to-bedside case study on tissue selective drug distribution. Bioorg Med Chem Lett 21, 2725–2731.

Pickett, K.E., and Wilkinson, R.G. (2015). Income inequality and health: a causal review. Social science & medicine 128, 316–326.

Pramod, A.B., Foster, J., Carvelli, L., and Henry, L.K. (2013). SLC6 transporters: Structure, function, regulation, disease association and therapeutics. Mol Aspects Med 34, 197–219.

Qiu, G.H., Xie, X., Xu, F., Shi, X., Wang, Y., and Deng, L. (2015). Distinctive pharmacological differences between liver cancer cell lines HepG2 and Hep3B. Cytotechnology 67, 1–12.

Reddy, V.S., Shlykov, M.A., Castillo, R., Sun, E.I., and Saier, M.H., Jr. (2012). The major facilitator superfamily (MFS) revisited. FEBS J 279, 2022–2035.

Rees, D.C., Johnson, E., and Lewinson, O. (2009). ABC transporters: the power to change. Nat Rev Mol Cell Biol 10, 218–227.

Reimer, R.J. (2013). SLC17: A functionally diverse family of organic anion transporters. Mol Aspects Med 34, 350–359.

Riniker, S., and Landrum, G.A. (2013). Open-source platform to benchmark fingerprints for ligand-based virtual screening. J Cheminform 5, 26.

Robinson, M.D., and Oshlack, A. (2010). A scaling normalization method for differential expression analysis of RNA-seq data. Genome Biol 11, R25.

Römpp, A., Guenther, S., Takats, Z., and Spengler, B. (2011). Mass spectrometry imaging with high resolution in mass and space (HR^2^ MSI) for reliable investigation of drug compound distributions on the cellular level. Anal Bioanal Chem 401, 65–73.

Sadras, V., and Bongiovanni, R. (2004). Use of Lorenz curves and Gini coefficients to assess yield inequality within paddocks. Field Crop Res 90, 303–310.

Santos, S.R., and Ochman, H. (2004). Identification and phylogenetic sorting of bacterial lineages with universally conserved genes and proteins. Environ Microbiol 6, 754–759.

Schlessinger, A., Matsson, P., Shima, J.E., Pieper, U., Yee, S.W., Kelly, L., Apeltsin, L., Stroud, R.M., Ferrin, T.E., Giacomini, K.M., et al. (2010). Comparison of human solute carriers. Protein Sci 19, 412–428.

Silver, N., Best, S., Jiang, J., and Thein, S.L. (2006). Selection of housekeeping genes for gene expression studies in human reticulocytes using real-time PCR. BMC molecular biology 7, 33.

Sontakke, S., Cadenas, M.B., Maggi, R.G., Diniz, P.P.V.P., and Breitschwerdt, E.B. (2009). Use of broad range 16S rDNA PCR in clinical microbiology. J Microbiol Meth 76, 217–225.

Sreedharan, S., Stephansson, O., Schiöth, H.B., and Fredriksson, R. (2011). Long evolutionary conservation and considerable tissue specificity of several atypical solute carrier transporters. Gene 478, 11–18.

Stanley, L.A., Horsburgh, B.C., Ross, J., Scheer, N., and Wolf, C.R. (2009). Drug transporters: Gatekeepers controlling access of xenobiotics to the cellular interior. Drug Metab Rev 41, 27–65.

Tatsumi, K., Ohashi, K., Taminishi, S., Okano, T., Yoshioka, A., and Shima, M. (2008). Reference gene selection for real-time RT-PCR in regenerating mouse livers. Biochem Biophys Res Commun 374, 106–110.

Taylor, P.M. (2014). Role of amino acid transporters in amino acid sensing. Am J Clin Nutr 99, 223S–230S.

Thul, P.J., Åkesson, L., Wiking, M., Mahdessian, D., Geladaki, A., Ait Blal, H., Alm, T., Asplund, A., Björk, L., Breckels, L.M., et al. (2017). A subcellular map of the human proteome. Science 356.

Torre, E., Dueck, H., Shaffer, S., Gospocic, J., Gupte, R., Bonasio, R., Kim, J., Murray, J., and Raj, A. (2017). A comparison between single cell RNA sequencing and single molecule RNA FISH for rare cell analysis. bioRxiv, 138289.

Tran, Q.N. (2011). Improving the Accuracy of Gene Expression Profile Classification with Lorenz Curves and Gini Ratios. Software Tools and Algorithms for Biological Systems 696, 83–90.

Tyzack, J.K., Wang, X., Belsham, G.J., and Proud, C.G. (2000). ABC50 interacts with eukaryotic initiation factor 2 and associates with the ribosome in an ATP-dependent manner. J Biol Chem 275, 34131–34139.

Uhlén, M., Fagerberg, L., Hallstrom, B.M., Lindskog, C., Oksvold, P., Mardinoglu, A., Sivertsson, Å., Kampf, C., Sjöstedt, E., Asplund, A., et al. (2015). Tissue-based map of the human proteome. Science 347, 1260419.

Vandesompele, J., De Preter, K., Pattyn, F., Poppe, B., Van Roy, N., De Paepe, A., and Speleman, F. (2002). Accurate normalization of real-time quantitative RT-PCR data by geometric averaging of multiple internal control genes. Genome Biol 3, RESEARCH0034.

Wang, F., Wang, J., Liu, D., and Su, Y. (2010). Normalizing genes for real-time polymerase chain reaction in epithelial and nonepithelial cells of mouse small intestine. Anal Biochem 399, 211–217.

Weidlich, I.E., and Filippov, I.V. (2016). Using the gini coefficient to measure the chemical diversity of small-molecule libraries. J Comput Chem 37, 2091–2097.

Wilkinson, R., and Pickett, K. (2009). The spirit level: why equality is better for everyone (London, Penguin Books).

Winter, G.E., Radic, B., Mayor-Ruiz, C., Blomen, V.A., Trefzer, C., Kandasamy, R.K., Huber, K.V.M., Gridling, M., Chen, D., Klampfl, T., et al. (2014). The solute carrier SLC35F2 enables YM155-mediated DNA damage toxicity. Nat Chem Biol 10, 768–773.

Wolbank, S., Stadler, G., Peterbauer, A., Gillich, A., Karbiener, M., Streubel, B., Wieser, M., Katinger, H., van Griensven, M., Redl, H., et al. (2009). Telomerase immortalized human amnion- and adipose-derived mesenchymal stem cells: maintenance of differentiation and immunomodulatory characteristics. Tissue Eng Part A 15, 1843–1854.

Wren, J.D. (2016). Bioinformatics programs are 31-fold over-represented among the highest impact scientific papers of the past two decades. Bioinformatics 32, 2686–2691.

Yarza, P., Yilmaz, P., Pruesse, E., Glöckner, F.O., Ludwig, W., Schleifer, K.H., Whitman, W.B., Euzéby, J., Amann, R., and Rosselló-Móra, R. (2014). Uniting the classification of cultured and uncultured bacteria and archaea using 16S rRNA gene sequences. Nat Rev Microbiol 12, 635–645.

Zampieri, M., Ciccarone, F., Guastafierro, T., Bacalini, M.G., Calabrese, R., Moreno-Villanueva, M., Reale, A., Chevanne, M., Burkle, A., and Caiafa, P. (2010). Validation of suitable internal control genes for expression studies in aging. Mech Ageing Dev 131, 89–95.

Zeigler, D.R. (2003). Gene sequences useful for predicting relatedness of whole genomes in bacteria. International journal of systematic and evolutionary microbiology 53, 1893–1900.

Zhuang, K., Vemuri, G.N., and Mahadevan, R. (2011). Economics of membrane occupancy and respiro-fermentation. Mol Syst Biol 7, 500.

